# Activation of Ca^2+^ phosphatase Calcineurin regulates Parkin translocation to mitochondria and mitophagy

**DOI:** 10.1101/2023.01.31.526442

**Authors:** Elena Marchesan, Alice Nardin, Sofia Mauri, Simone Di Paola, Monica Chinellato, Sophia von Stockum, Joy Chakraborty, Stephanie Herkenne, Valentina Basso, Emilie Schrepfer, Oriano Marin, Laura Cendron, Diego L. Medina, Luca Scorrano, Elena Ziviani

**Affiliations:** Department of Biology, University of Padova, Padova, Italy; Medical Research Council Protein Phosphorylation and Ubiquitylation Unit, University of Dundee, Dundee, UK; Telethon Institute of Genetics and Medicine (TIGEM), Pozzuoli, Naples, Italy; Institute for Experimental Endocrinology and Oncology (IEOS), National Research Council (CNR), Napoli, Italy; Dulbecco-Telethon Institute, Venetian Institute of Molecular Medicine (VIMM), Padova, Italy; Department of Biomedical Sciences (DSB), University of Padova, Padova, Italy; Medical Genetics Unit, Department of Medical and Translational Science, Federico II University, Naples, Italy

## Abstract

Selective removal of dysfunctional mitochondria via autophagy is crucial for the maintenance of cellular homeostasis. This event is initiated by the translocation of the E3 ubiquitin ligase Parkin to damaged mitochondria, and it requires the Serine/Threonine-protein kinase PINK1. In a coordinated set of events, PINK1 operates upstream of Parkin in a linear pathway that leads to the phosphorylation of Parkin, Ubiquitin, and Parkin mitochondrial substrates, to promote ubiquitination of outer mitochondrial membrane proteins. Ubiquitin decorated mitochondria are selectively recruiting autophagy receptors,which are required to terminate the organelle via autophagy. In this work we show a previously uncharacterized molecular pathway that correlates the activation of the Ca^2+^-dependent phosphatase Calcineurin to Parkin-dependent mitophagy. Calcineurin downregulation or genetic inhibition prevents Parkin translocation to CCCP-treated mitochondria, and impairs stress-induced mitophagy, whereas Calcineurin activation promotes Parkin mitochondrial recruitment and basal mitophagy. Calcineurin interacts with Parkin, and promotes Parkin translocation in the absence of PINK1, but requires PINK1 expression to execute mitophagy in MEF cells. Genetic activation of Calcineurin *in vivo* boosts basal mitophagy in neurons, and corrects locomotor dysfunction and mitochondrial respiratory defects of a *Drosophila* model of impaired mitochondrial functions.

Our study identifies Calcineurin as a novel key player in the regulation of Parkin translocation and mitophagy.

## Introduction

The Ubiquitin Proteasome System (UPS) and mitophagy are dysregulated in many neurological diseases, including Parkinson’s Disease (PD), a neurodegenerative condition characterized by dopaminergic neuron loss^1,2^, accumulation of ubiquitinated unfolded protein aggregates^3,4^, mitochondrial dysfunction and mitophagy dysregulation^5,6^. Although most PD cases are sporadic, a small proportion derives from mutations in PD associated genes^7,8^, which have been identified by characterizing familiar Mendelian inherited PD forms, and includes mutations in the mitophagy genes PINK1 and Parkin^9–11^. Parkin belongs to the RBR (RING-between-RING) type of E3 ubiquitin ligases^12^, also known as RING/HECT hybrids, consisting of an ubiquitin like domain (Ubl), followed by two RING fingers domains (RING0 and RING1), an in between RING finger domain (IBR), a linker domain called Repressor Element of Parkin (REP) and a third RING finger domain called RING2^13–16^. Under basal conditions, Parkin activity is repressed and the protein maintains a close structure with the Ubl domain and the REP fragment occluding the RING1 domain, and the RING0 domain impeding on the catalytic site-containing RING2 domain^14–16^. The gene product has a number of neuroprotective roles and pleiotropic functions and its activation is involved in many different survival pathways^17–19^, including those that are affecting mitochondrial function by regulating mitochondria quality control^20–24^. How Parkin controls so many different cellular processes is under intense investigation. However, by being a versatile E3 ubiquitin ligase that both promotes degradative Lys 48-mediated ubiquitination and nonclassical, proteosomal-independent ubiquitination^25–31^, Parkin has the potential of controlling a broad subset of cellular processes. Not surprisingly, Parkin activity is repressed under basal conditions, and its activation is tightly regulated by a number of molecular processes, which are largely mediated by post translational modifications^13,14,16^. Parkin function is closely related to the activity of another PD-related gene, PARK6, which encodes for a protein called PINK1^32–34^. PINK1 is a Serine/Threonine kinase that is imported into mitochondria, where it gets cleaved by the inner membrane protease PARL and then eliminated by the proteasome^35–37^. On depolarized mitochondria, PINK1 accumulates on the outer mitochondrial membrane (OMM), where it promotes Parkin translocation and mitochondrial recruitment^33,34,38,39^. Independent studies showed that PINK1-mediated phosphorylation of Parkin and Ubiquitin at residue Serine 65 (Ser65) is required for Parkin translocation to defective mitochondria, and for its E3-ubiquitin ligase activity^40–43^. This process leads to Parkin-dependent ubiquitination and proteasomal degradation of OMM proteins, and to the selective autophagy of damaged mitochondria^44–48^. Moreover, PINK1 phosphorylates a number of Parkin substrates including the pro-fusion protein Mitofusin, Mfn2, which work as Parkin receptor^49^. It was proposed that Parkin-dependent ubiquitination of Mitofusins prevents mitochondrial fusion of dysfunctional mitochondria by promoting proteasomal degradation of Mfn1 and Mfn2^24^, and segregates depolarized mitochondria from the mitochondrial network, impairing their ability to refuse^50^. In addition, PINK1 directly promotes mitochondrial fission by regulating recruitment and activation of pro-fission protein Drp1^51^. In resting conditions, Drp1 is phospho-inhibited at Ser 637 by AKAP1-PKA (A-kinase anchoring protein complex and protein kinase A, respectively)^52^. Following mitochondria damage, PINK1 becomes active and disrupts the AKAP1-PKA complex to promote Drp1-dependent fission^51^. Moreover, PINK1 directly phosphorylates Drp1 on Ser 616 to regulate mitochondrial fission^53^. Interestingly, Drp1 is selectively recruited to dysfunctional mitochondria in the proximity of PINK1/Parkin suggesting that mitochondrial division occurs at sites where the PINK1/Parkin-dependent mitochondrial clearance program is initiated^54^. Expression of dominant-negative Drp1 to inhibit mitochondrial fission prevents mitophagy, indicating the importance of fission in mitophagy^55^, which is also supported by studies on yeasts^56^. Translocation of Drp1 to mitochondria is mediated by selective dephosphorylation of residue Serine 637, which is controlled by Ca^2+^ and Ca^2+^/Calmodulin dependent phosphatase Calcineurin (CaN)^52^. Rise of cytosolic Ca^2+^ associated to mitochondrial depolarization^57^, leads to CaN-dependent Drp1 dephosphorylation and Drp1 mitochondrial recruitment to promote mitochondrial fission^52^, which is required to execute mitophagy^58,59^. CaN also dephosphorylates transcription factor TFEB to promote its nuclear translocation and the expression of autophagy and lysosomal genes to induce autophagy and lysosomal biogenesis^60^.

In this work, we show that Ca^2+^/Calmodulin phosphatase CaN is required for Parkin mitochondrial recruitment and mitophagy in CCCP-treated mouse embryonic fibroblast cells (MEFs). CaN activation is sufficient to promote Parkin translocation under basal condition in a PINK1-independent fashion. Genetic activation of CaN *in vivo* promotes basal mitophagy in neurons, and corrects locomotor dysfunction and mitochondrial defects of a *Drosophila* model of impaired mitochondrial function.

## Results

### Parkin translocation to mitochondria is regulated by Calcineurin

We transfected mouse embryonic fibroblasts (MEFs),, with fluorescent mCherry-Parkin and mitochondrial targeted YFP (mitoYFP), and analyzed Parkin subcellular localization by confocal microscopy, following established experimental protocols ^34,38,46^. Consistent with previous studies^33,38,46,61^, we observed that mCherry-Parkin was predominantly located in the cytosol in non-treated cells (Figure 1A, upper panel). Following the treatment with uncoupling agent carbonyl cyanide m-chlorophenylhydrazone (CCCP), a significant proportion of cells (75,7±1,9%) showed mCherry-Parkin accumulated on or near fragmented mitochondria (Figure 1A, lower panel), forming discrete dots, which we called puncta (quantified in Figure 1B). An automated analysis of the confocal images with Squassh, an ImageJ plugin that calculates the degree of colocalization between two channels^62^, allowed to further consolidate this result. In this analysis, the colocalization coefficient computed by the Squassh plugin ranges from 0 to 1, where 0 indicates no colocalization, and 1 perfect colocalization between mCherry-Parkin and YFP labeled mitochondria. According to this analysis, Squassh index raised from 0,29±0,03 to 0,62±0,06 following CCCP treatment, indicative of increased Parkin mitochondrial recruitment (Figure 1C). A small proportion of Parkin puncta did not seem to colocalize with mitochondria (Figure 1A, arrowhead) suggesting a potential recruitment of Parkin to other organelles. To test this hypothesis, we cotransfected mCherry-Parkin expressing MEFs with the lysosomal marker LAMP1GFP or the endosomal marker Rab5BGFP. Colocalization analysis using Squassh demonstrated that in CCCP-treated cells a small proportion of Parkin puncta colocalized with lysosomes (Figure 1D, quantified in Figure 1E), whether no significant recruitment to endosomes occurs (Figure 1F; quantified in Figure 1G). The addition of the protophore CCCP, required for triggering Parkin translocation, induces a transient increase of Ca^2+^ influx^57^. Importantly, Ca^2+^ chelation with BAPTA abolishes Parkin translocation (Supplementary Figure 1), indicating a role for Ca^2+^-dependent signal in the regulation of Parkin mitochondrial recruitment. Because mitochondria need to fragment to be engulfed by the autophagosome^58,59^, and rise of cytosolic Ca^2+^ associated to mitochondrial depolarization leads to Calcineurin (CaN)-dependent Drp1 fission^52^ and autophagy^60^, it is conceivable that CaN may also play a role in Parkin recruitment and Parkin-dependent mitophagy. Treatment with FK506, an immunosuppressive agent that blocks CaN without affecting the permeability transition pore (PTP)^63^, impaired Parkin translocation (Supplementary Figure 2). This was also the case upon addition of smaller concentration of CCCP, and following treatment with other mitochondrial damaging compounds that promoted mitochondrial depolarization (Supplementary Figure 3). To further support this result, we took advantage of the existing dominant negative mutant of CaN (ΔCnA^H151Q^)^64,65^. CaN is a heterodimer, composed of a catalytic subunit (CnA) that binds calmodulin and a regulatory subunit (CnB) that binds Ca^2+^. Ca^2+^/calmodulin activates CaN upon binding to the calmodulin-binding domain of CnA and inducing the dissociation of the autoinhibitory domain from the catalytic domain^52^. ΔCnA^H151Q^ dominant negative mutant misses the calmodulin binding domain and the autoinhibitory domain and harbors an inactivating His-151 to Gln point mutation. We cotransfected MEFs with mCherry-Parkin and dominant negative CaN (CnB plus ΔCnA^H151Q^) and looked at Parkin localization. As previously showed, CCCP-induced Parkin mitochondrial recruitment was clearly visible following 3hrs CCCP (Figure 2A, left panel). This event was completely abolished when in presence of the dominant negative CaN, ΔCnA^H151Q^ (Figure 2A, right panel; quantified in Figure 2B-C). CaN downregulating cells (Supplementary Figure 4) also exhibited a significant decrease in Parkin recruitment upon CCCP treatment (Figure 2D, quantified in Figure 2E-F).

**Figure 1:**
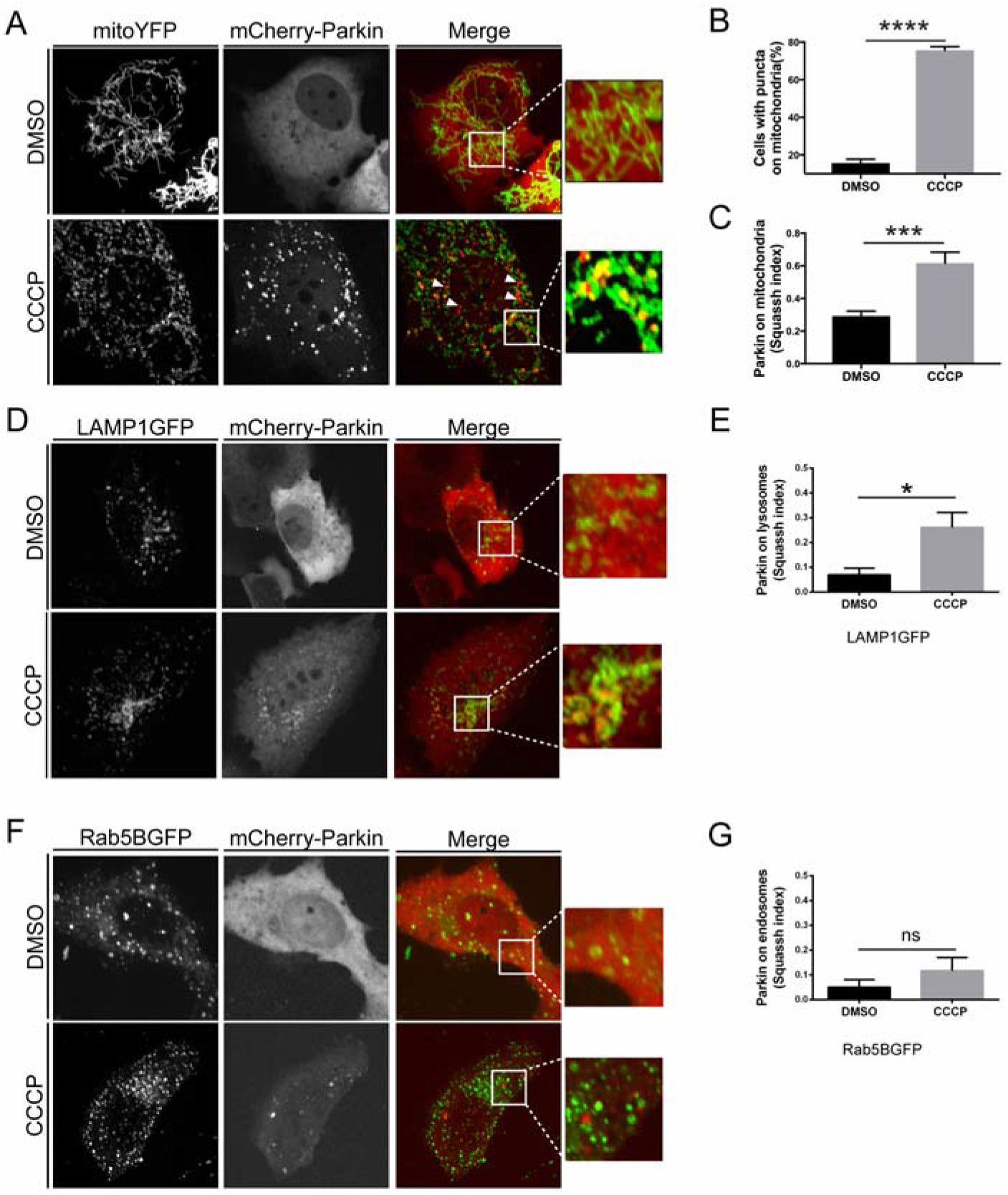
Parkin translocation to mitochondria is regulated by Calcineurin. (A) Representative confocal images of MEF cells transfected with mCherry-Parkin and mito-YFP for 2 days before being treated with DMSO or 10 μM CCCP for 3 hours. The panels on the right show enlarged merged views of the boxed areas (B) Quantification of A. Graph bar shows mean±SEM of percentage of cells with mCherry-Parkin on mitochondria for at least ≥ 300 cells per biological replicate. Student’s t-test (n=9-10; p<0.0001). (C) Quantification of A by using Squassh. The graph bars show mean±SEM of Squassh colocalization coefficient for at least ≥ 50 images per biological replicate. 0=no colocalization, 1=perfect colocalization. Student’s t-test (n=5; p<0.0001) (D) Representative confocal images of MEF cells transfected with mCherry-Parkin and LAMP1GFP for 2 days and treated with 10 μM CCCP for 3 hours. (E) Quantification of D by using Squassh. The graph bars show mean ± SEM of Squassh colocalization coefficient for at least ≥ 50 images per biological replicate. Student’s t-test (n=3; p<0.05) (F) Representative confocal images of MEF cells transfected with mCherry-Parkin and with Rab5GFP for 2 days and treated with 10 μM CCCP for 3 hours (G) Quantification of F by using Squassh. The graph bars show mean ± SEM of Squassh colocalization coefficient for at least ≥ 50 images per biological replicate. Student’s t-test (n=3; p<0.05).

**Figure 2:**
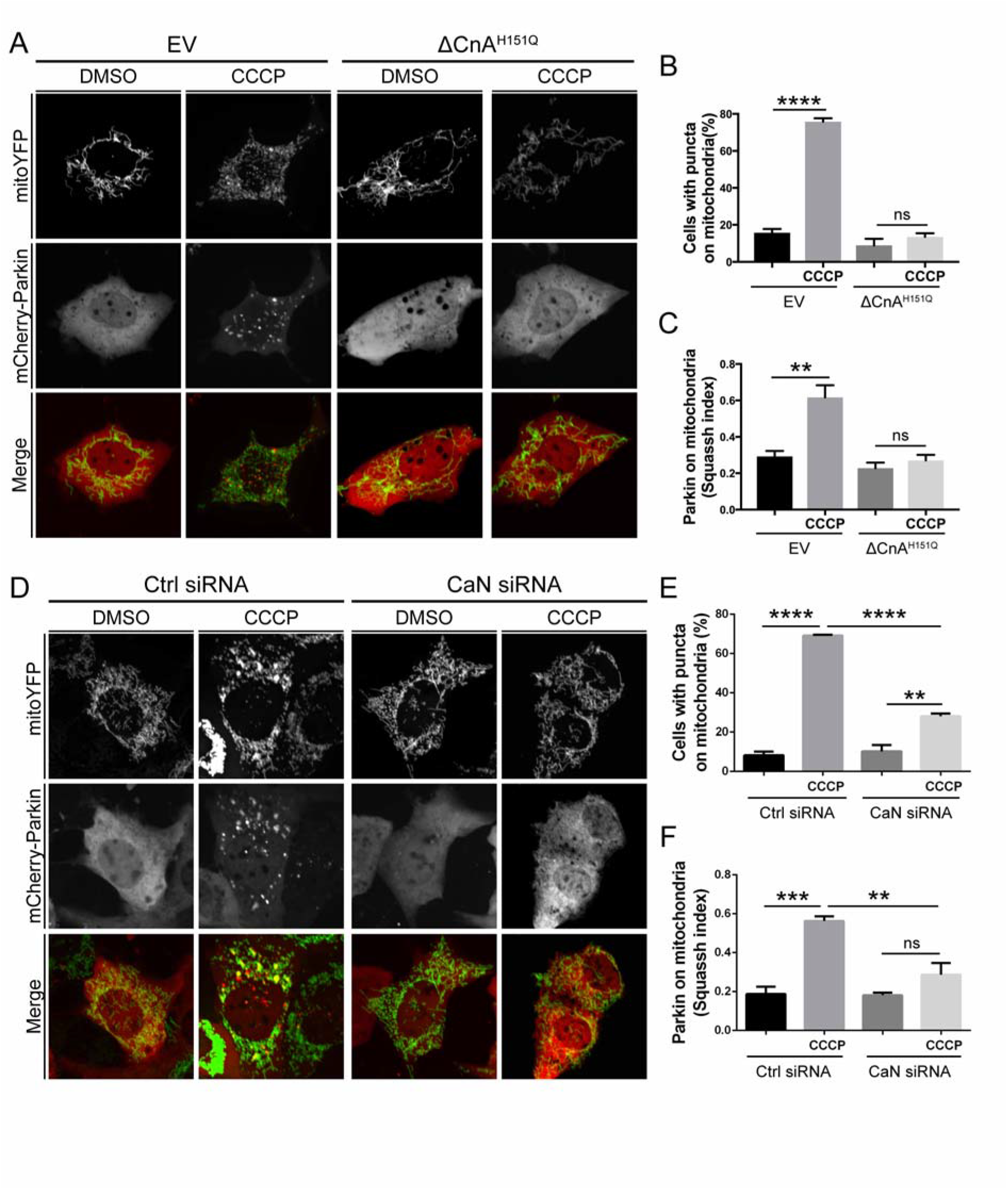
Parkin translocation to mitochondria is regulated by Calcineurin. (A) Representative confocal images of MEF cells transfected with mCherry-Parkin, mito-YFP and dominant negative CaN (ΔCnA^H151Q^) or the empty vector (EV) for 2 days before being treated with DMSO or 10 μM CCCP for 3 hours. (B) Quantification of (A). Graph bar shows mean ± SEM of percentage of cells with mCherry-Parkin on mitochondria for at least ≥ 300 cells per biological replicate. Two-way ANOVA followed by Tukey’s multiple comparison test (n=3-9; p<0.0001). (C) Quantification of (A) using Squassh. The graph bars show mean±SEM of Squassh colocalization coefficient for at least ≥ 50 images per biological replicate. 0=no colocalization, 1=perfect colocalization. Two-way ANOVA followed by Tukey’s multiple comparison test (n=4-5; p<0.001) (D) Representative confocal images of MEF cells transfected with mCherry-Parkin, mito-YFP in which CaN was downregulated, and relative control. (E) Quantification of (D). Graph bar shows mean ± SEM of percentage of cells with mCherry-Parkin on mitochondria for at least ≥300 cells per biological replicate. Two-way ANOVA followed by Tukey’s multiple comparison test (n=3; p<0.01). (F) Quantification of (D) using Squassh. The graph bar shows mean ± SEM of Squassh colocalization coefficient for at least ≥ 50 images per biological replicate. 0=no colocalization, 1=perfect colocalization. At least 3 independent experiments were performed. Two-way ANOVA followed by Tukey’s multiple comparison test (n=3; p<0.01)

### Parkin translocation is induced by Calcineurin in the absence of PINK1

Parkin translocation and activity are strictly controlled by mechanisms of autoinhibition that can be released by PINK1^13,15,16,66–68^. The current model for the mechanism of Parkin activation and recruitment by PINK1 depicts that under basal conditions Parkin activity is repressed and the protein maintains a close structure with the ubiquitin-like domain (UBL) and the repressor element of Parkin (REP) fragment occluding the RING1 domain and the RING0 domain impeding on the catalytic site-containing RING2 domain^14–16^. Structural studies have provided evidence that phosphorylation of Ubiquitin and Parkin at Serine 65 by PINK1 causes displacement of the inhibitory UBL and stretches the REP^43^. This affects Parkin autoinhibitory structure and contributes to promote Parkin conformational change that leads to Parkin mitochondrial recruitment and activation^43,69,70^. In line with this model and as already reported^38,39^, we found that Parkin does not translocate to CCCP-treated mitochondria in the absence of PINK1 (Figure 3A, left panel). Intriguingly, expression of constitutive active CaN (CnB plus ΔCnA) promoted Parkin translocation in PINK1 KO cells, even in the absence of CCCP (Figure 3A, right panel; quantified in Figure 3B-C), a condition that was hold true also in PINK1 wild-type cells (Supplementary Figure 5). Moreover, in PINK1 KO MEFs transfected with phospho-mimetic Ub (Ub S65E), a large proportion of phospho-mimetic mCherry-Parkin (Parkin S65E) translocated to CCCP-treated mitochondria (Figure 3D, left panel), a condition that was abolished upon expression of CaN dominant negative ΔCnA^H151Q^ (Figure 3D, right panel; quantified in Figure 3E-F). Based on these results, it is expected that expression of CaN can affect Parkin conformational change in the absence of PINK1 to promote Parkin translocation. Because conformational changes are paralleled by changes in protein solubility, which can be assessed by thermal shift^71,72^, we performed a thermal stability assay for Parkin to test this hypothesis. Expression of constitutive active CaN leads to a decrease in Parkin thermal stability in wild type (Figure 3G-H) and PINK1 KO cells (Figure 3I-J), supporting the hypothesis that Parkin undergoes a conformational change in this condition regardless PINK1 expression.

**Figure 3:**
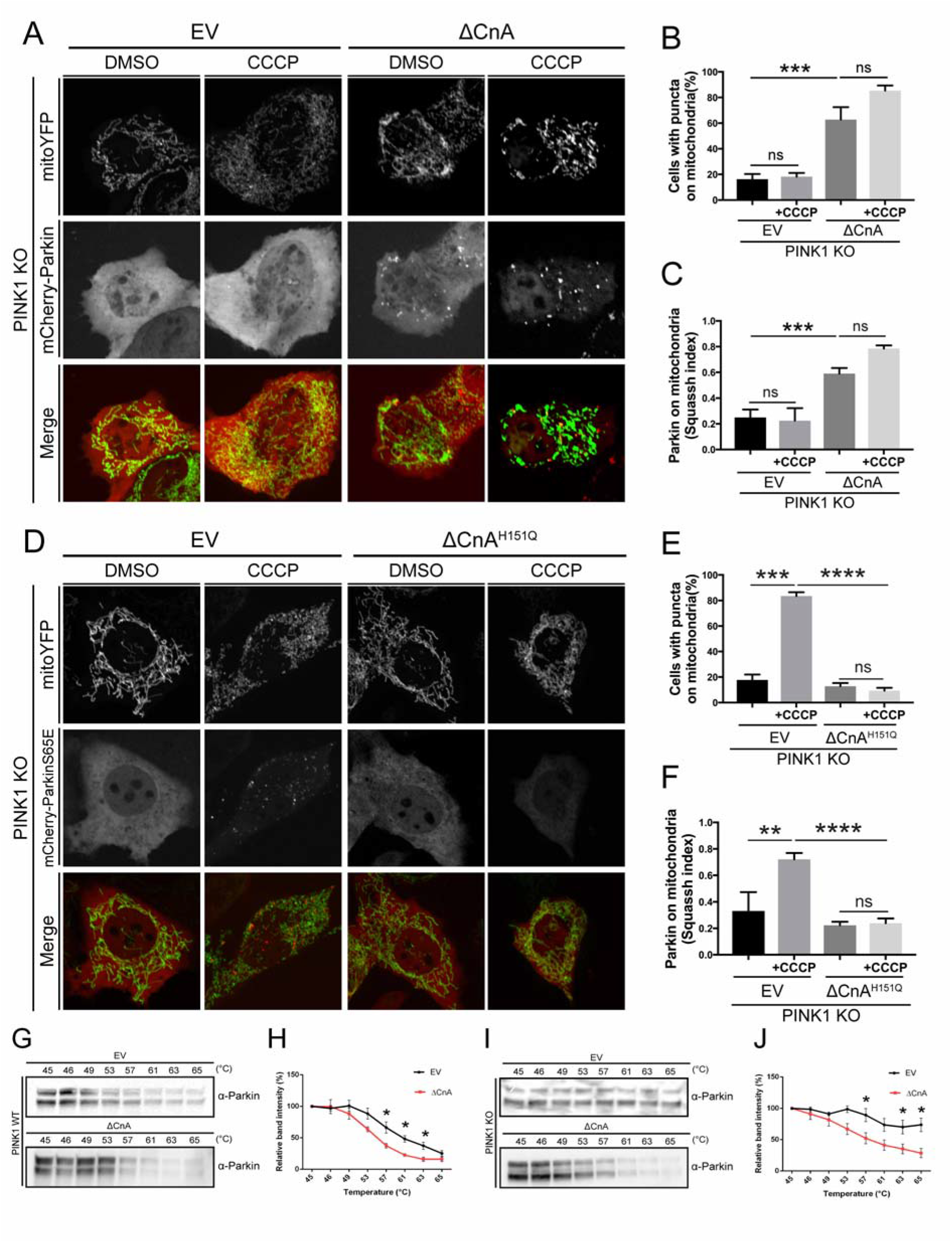
Parkin translocation is induced by Calcineurin in the absence of PINK1. (A) Representative confocal images of PINK1 KO MEF cells transfected with mCherry-Parkin, mito-YFP and with empty vector (EV) or constitutively active CaN (ΔCnA). (B) Quantification of (A) Graph bar shows mean ± SEM of percentage of cells with mCherry-Parkin on mitochondria for at least ≥ 300 cells per biological replicate. Two-way ANOVA followed by Tukey’s multiple comparison test (n=3-4; p<0.001). (C) Quantification of (A) using Squassh. The graph bars show mean±SEM of Squassh colocalization coefficient for at least ≥ 50 images per biological replicate. 0=no colocalization, 1=perfect colocalization. At least 3 independent experiments were performed. Two-way ANOVA followed by Tukey’s multiple comparison test (n=3-4; p<0.001). (D) Representative confocal images of PINK1 KO MEF cells transfected with mCherry-ParkinS65E, UbS65E, mito-YFP and with empty vector (EV) or dominant negative CaN (ΔCnA^H151Q^) (E) Quantification of (D) Graph bar shows mean ± SEM of percentage of cells with mCherry-Parkin on mitochondria for at least ≥ 300 cells per biological replicate. Two-way ANOVA followed by Tukey’s multiple comparison test (n=4; p<0.0001). (F) Quantification of (D) using Squassh. The graph bars show mean±SEM of Squassh colocalization coefficient for at least ≥ 50 images per biological replicate. 0=no colocalization, 1=perfect colocalization. Two-way ANOVA followed by Tukey’s multiple comparison test (n=4; p<0.0001). (G) Parkin thermal stability assay. WT MEFs expressing constitutive active CaN (ΔCnA) or empty vector (EV) were suspended in PBS and snap-freezed in liquid nitrogen before being aliquoted into a PCR strip and incubated at the indicated temperature for 3 min. The lysates were centrifugated at high speed and the soluble fraction was loaded into SDS-PAGE gel. Representative Western blotting analysis for Parkin stability is shown. (H) Densitometric analysis of (G). Student’s t-test (n=4; p<0.05). (I) Parkin thermal stability assay. PINK1 KO MEFs expressing constitutive active CaN (ΔCnA) or empty vector (EV) were suspended in PBS and snap-freezed in liquid nitrogen before being aliquoted into a PCR strip and incubated at the indicated temperature for 3 min. The lysates were centrifugated at high speed and the soluble fraction was loaded into SDS-PAGE gel. Representative Western blotting analysis for Parkin stability is shown. (J) Densitometric analysis of (I). Student’s t-test (n=6; p<0.05).

Thus, expression of constitutively active CaN promotes Parkin recruitment in PINK1 KO cells that is in the absence of PINK1-dependent phosphorylation of Parkin and Ubiquitin.

### Calcineurin interacts with Parkin

Our results prompted us to evaluate the possibility of an interaction between CaN and Parkin. To evaluate this possibility, we first performed an *in vitro* interaction assay by generating affinity-purified recombinant His-tagged Parkin from bacteria, which was coupled to a His-affinity resin and incubated with protein lysate extracted from cells expressing Flag-tagged CaN. The resin was washed to remove nonspecifically adhering proteins, and Laemmli buffer was used to elute the complexes from the resin (Figure 4A). Importantly, in the eluted complexes we were able to identify CaN (Figure 4B), indicating that CaN binds to Parkin *in vitro*. The interaction was specific for CaN because an unrelated Flag-tagged protein that is not supposed to interact with Parkin (USP14-Flag) was not retrieved in the eluted complexes when protein lysate from cells expressing USP14-Flag was incubated with the resin. Likewise, no interaction was retrieved when His-tagged MEF2D, which does not interact with CaN, was used as a bait to retrieve CaN (Figure 4B).

**Figure 4:**
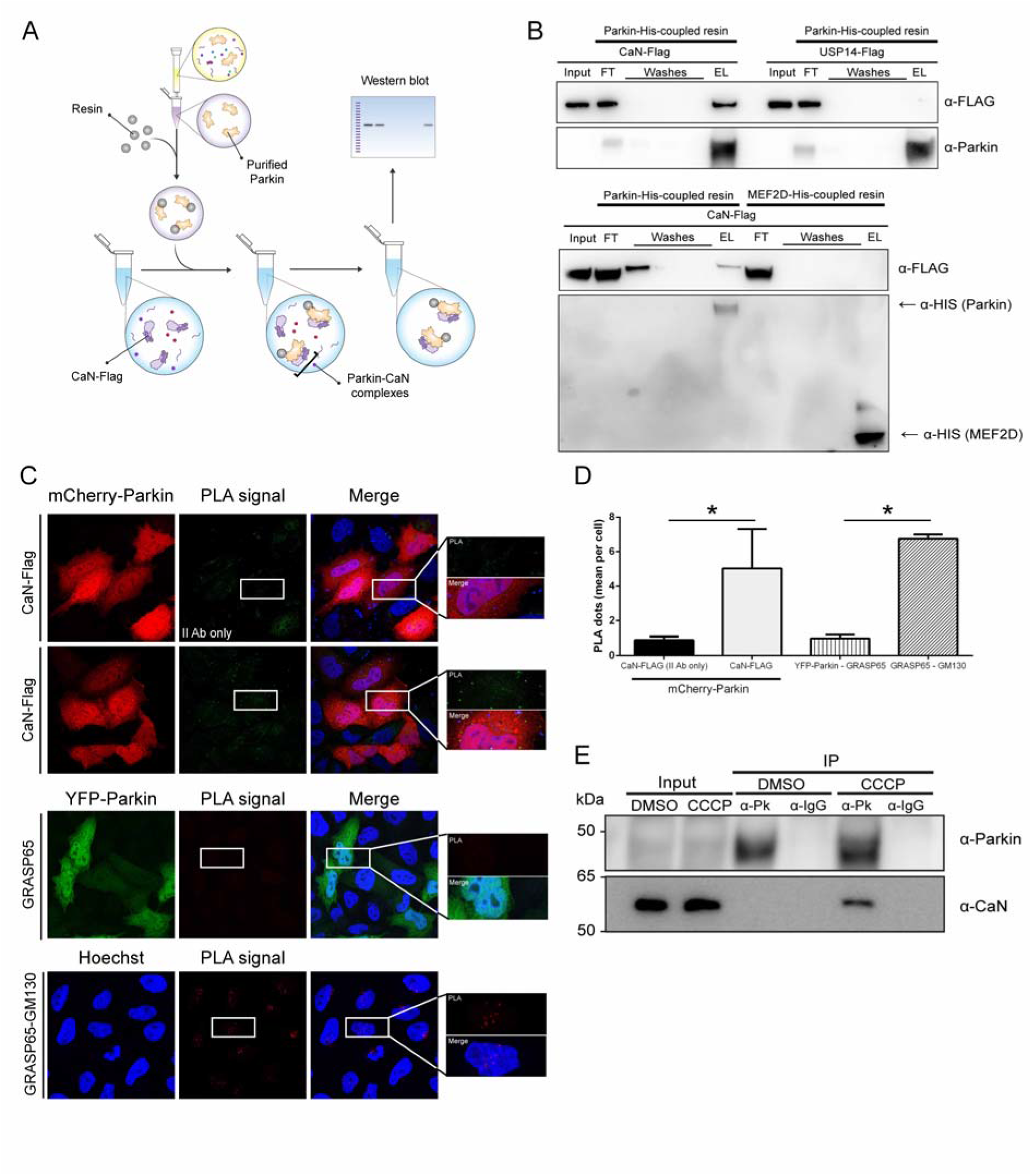
Calcineurin interacts with Parkin. (A) *In vitro* interaction assay: schematic representation. Affinity-purified recombinant His-tagged Parkin is produced from bacteria, and coupled to a His-affinity resin to generate Parkin-His-resin. The resin is incubated with protein lysate extracted from cells expressing Flag-tagged CaN. The resin is washed to remove nonspecifically adhering proteins, and Imidazole is used to elute the complexes from the resin. The obtained eluate is separated by SDS-PAGE and analyzed by immunoblotting. (B) *In vitro* interaction assay. His-tagged Parkin or His-tagged MEF2D coupled to a His-affinity resin are incubated with cell lysate obtained from MEFs transfected with CaN-Flag or USP14-Flag, as indicated. The eluate is separated by SDS-PAGE and analyzed by immunoblotting using anti-Flag, anti-Parkin and anti-His antibodies (FT=flow through; EL=eluate). (C) Representative images of Proximity Ligation Assay (PLA) performed on HeLa cells transfected with Parkin-mCherry and CaN-FLAG (PPP3CB-FLAG). After fixation, cells were processed for investigating Parkin proximity interactions by incubating with anti-Parkin antibody and anti-PPP3CB antibody, and corresponding secondary antibodies, or secondary antibodies alone as negative control. CaN and Parkin interactions are represented by green dots. As negative biological control for PLA we used anti-Parkin antibody and anti-GRASP65 antibody in YPF-Parkin expressing HeLa cells. As positive control for PLA, we used anti-GRASP65 antibody and anti-GM130 antibody. GRASP65 is a peripheral membrane protein that resides in the *cis-*Golgi apparatus, and interacts with GM130 (also expressed in the *cis*-Golgi network) but not with Parkin. GRASP65 and GM130 interactions are represented by red dots. White squares contain higher-magnification images. (D) Quantification of (C). Graph bar shows mean ± SEM of PLA dots per cell for at least ≥ 350 cells per biological replicate. Student’s t-test (n=3-4; p<0.05) (E) HEK 293T cells were treated with DMSO or CCCP-2hrs and subjected to immunoprecipitation (IP) of Parkin using mouse anti-Parkin antibody or anti-mouse IgG as negative control. Western Blot analysis was performed with rabbit anti-CaN antibody or mouse anti-Parkin antibody on the pulled down samples. Inputs represent 5% of the protein lysates and IP eluate 100% of the protein lysates. Mouse IgG TrueBlot^®^ ULTRA enabled detection of Parkin band, without hindrance by interfering immunoprecipitating IgG heavy chains.

We next performed a proximity ligation assay (PLA) to investigate Parkin and CaN interaction *in situ*. We were able to visualize discrete spots (PLA signal) in HeLa cells co-expressing mCherry-Parkin and CaN-Flag, representative of Parkin-CaN close proximity. In this assay, the interaction between GRASP65 and Parkin was used as negative biological control, whereas the interaction between GRASP65 and GM130 was used as positive control for PLA^73,74^. PLA detection dots indicate positive interaction (Figure 4C; quantified in Figure 4D). The specificities of the antibodies used in the PLA were tested by immunofluorescence (Supplementary Figure 6).

Finally, to evaluate the possibility of an interaction at the endogenous level, we performed an immunoprecipitation (IP) assay in HEK 293T cells, which express relatively high levels of endogenous Parkin. HEK cells were treated with DMSO or CCCP for 2hrs, and endogenous Parkin from cell lysate was captured by specific Parkin antibody. The antibody-protein complexes were pulled out of the sample using Protein A-coupled agarose beads. Endogenous CaN co-immunoprecipitated with Parkin in these complexes, indicating that endogenous CaN and Parkin interact. Notably, the interaction between endogenous Parkin and CaN was detectable only upon CCCP treatment to activate CaN, and not under basal conditions (Figure 4E). To further strengthen this result, we also performed the reverse IP, i.e. we pulled down endogenous CaN from lysate of cells treated with CCCP as before, and we were able to retrieve endogenous Parkin from the pulled down sample (Supplementary Figure 7).

Thus, CaN interacts with Parkin.

### Parkin translocation is induced by Calcineurin independently of Drp1 mitochondrial recruitment and activity

Previous reports demonstrated that CaN regulates the phosphorylation status of mitochondrial pro fission protein Drp1, and that dephosphorylation by CaN on Serine 637 is required for Drp1 translocation and in the process of mitochondrial fission^52^. Because CaN activation promotes mitochondrial recruitment of Drp1 and Parkin, we addressed whether these two events were correlated. Interestingly, expression of constitutive dephosphorylated Drp1 (Drp1 S637A) that was shown to be mostly mitochondrial and to promote mitochondrial fragmentation^52^, did not affect Parkin translocation in presence of CaN dominant negative (Figure 5A-B). Moreover, promotion of Parkin recruitment by expression of constitutive active CaN was not inhibited in presence of Drp1 dominant negative (Drp1 K38A) (Figure 5C-D), indicating that Parkin translocation is independent of Drp1 recruitment to the mitochondria to drive mitochondrial fragmentation.

**Figure 5:**
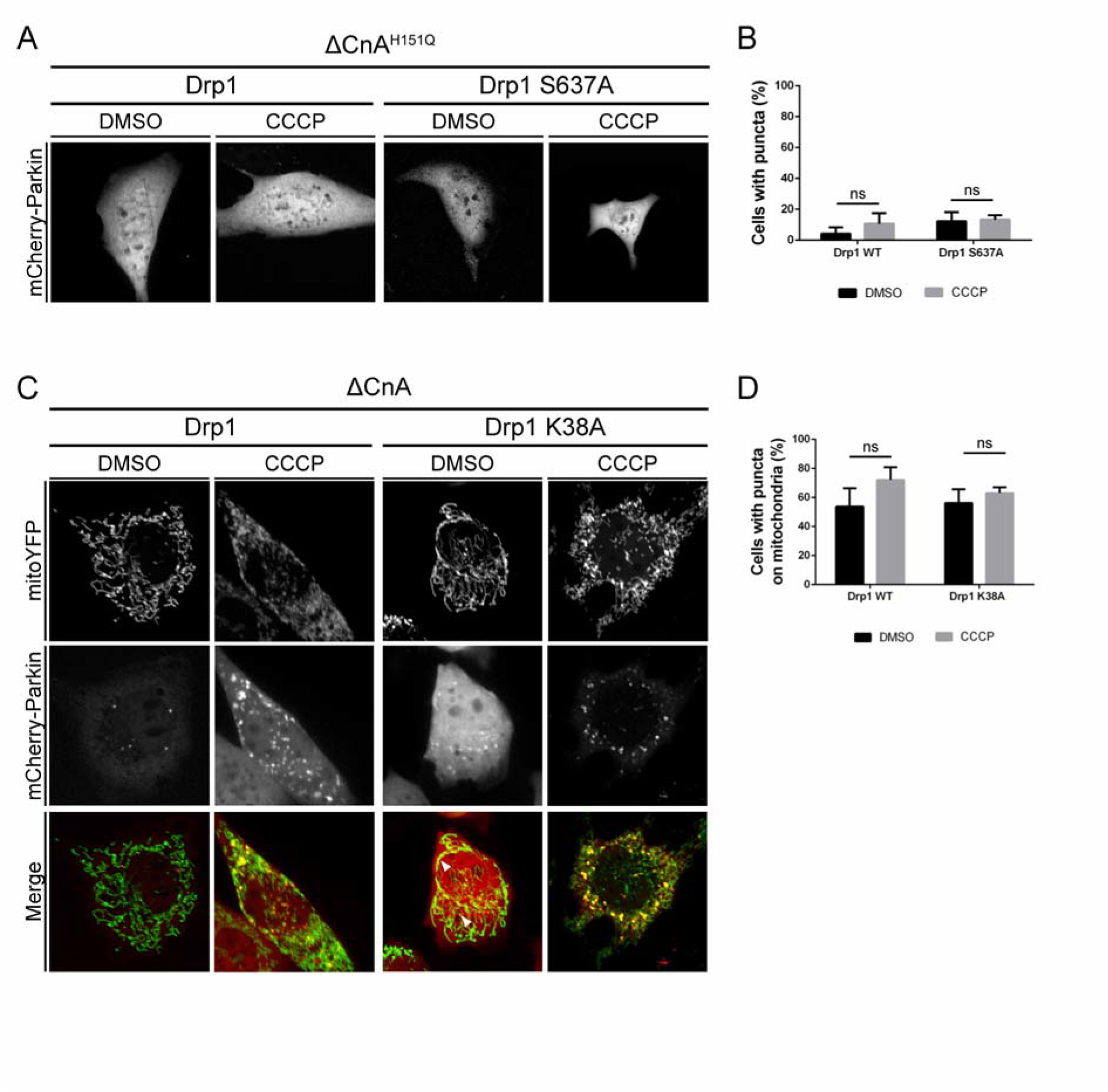
Parkin translocation is induced by Calcineurin independently of Drp1 mitochondrial recruitment and activity. (A) Representative confocal images of wild type MEF cells transfected with mCherry-Parkin and YFP-Drp1 or YFP-Drp1 S637A plus dominant negative CaN (ΔCnA^H151Q^) for 2 days before being treated with DMSO or 10 μM CCCP for 3 hours. (B) Quantification of A. Graph bar shows mean ± SEM of percentage of cells with mCherry-Parkin puncta for ≥ 300 cells per biological replicate. Two-way ANOVA followed by Tukey’s multiple comparison test (n=3). (C) Representative confocal images of wild type MEF cells transfected with mCherry-Parkin, mito-YFP and constitutive active CaN (ΔCnA) plus Drp1 or Drp1 K38A for 2 days before being treated with DMSO or 10 μM CCCP for 3 hours. (D) Quantification of C. Graph bar shows mean ± SEM of percentage of cells with mCherry-Parkin on mitochondria for ≥ 300 cells per biological replicate. Two-way ANOVA followed by Tukey’s multiple comparison test (n=3;).

### Calcineurin is required for CCCP-induced mitophagy

Our data demonstrate that CaN interacts with Parkin, and that CaN activity is required for Parkin recruitment to mitochondria. What about mitophagy? We used four different approaches to evaluate mitophagy *in vitro* in MEFs, namely (i) quantification of mitochondrial mass, estimated by western blotting analysis of inner mitochondrial membrane protein ATP synthase (ATP5A), (ii) confocal analysis of LC3-decorated mitochondria, (iii) FACS and confocal analysis of mitochondria-targeted fluorescent probe mt-Keima, and (iv) electron microscopy (EM).

Because MEFs have negligible levels of Parkin^44,75–77^, which might affect our ability to investigate Parkin-dependent events, we utilized a well established experimental system^34,46,78–80^ in which we introduced exogenous Parkin into MEFs by generating a Parkin-flag stable cell line by retroviral infection^81^. MEFs stably expressing Parkin-flag were transfected with dominant negative CaN (ΔCnA^H151Q^) or corresponding empty vectors, and mitochondrial mass was assessed following treatment with CCCP. In this condition, mitochondrial protein ATP synthase subunit alpha, ATP5A was lost after 36 hrs CCCP treatment, while it was retained in cells expressing dominant negative CaN, ΔCnA^H151Q^ (Figure 6A-B). Cells downregulating CaN also exhibited impaired CCCP-induced mitochondrial degradation (Figure 6C-D). These quantitative immunoblotting data were confirmed by confocal analysis of colocalization of mito-Kate labelled mitochondria with GFP-LC3 labelled autophagosomes. Cells expressing dominant negative CaN (Figure 6E) or downregulating CaN (Figure 6F) presented impaired LC3-mitochondria colocalization, following CCCP (quantified in Figure 6G-H). To further evaluate mitophagy in our model system, we infected Parkin expressing MEFs with mt-Keima, a pH-dependent fluorescence probe targeted to the mitochondrial matrix, which has different excitation spectra at neutral and acidic pH ^82^. Keima has a single emission peak at 620 nm with a bimodal excitation spectrum. These properties of mt-Keima allow rapid FACS determination of “acidic” mitochondria undergoing autolysosome degradation. The mt-Keima assay showed that CCCP-induced mitophagy was greatly reduced in cells downregulating CaN, in line with the results obtained with the biochemical approaches (Figure 6I and Supplementary Figure 8).

**Figure 6:**
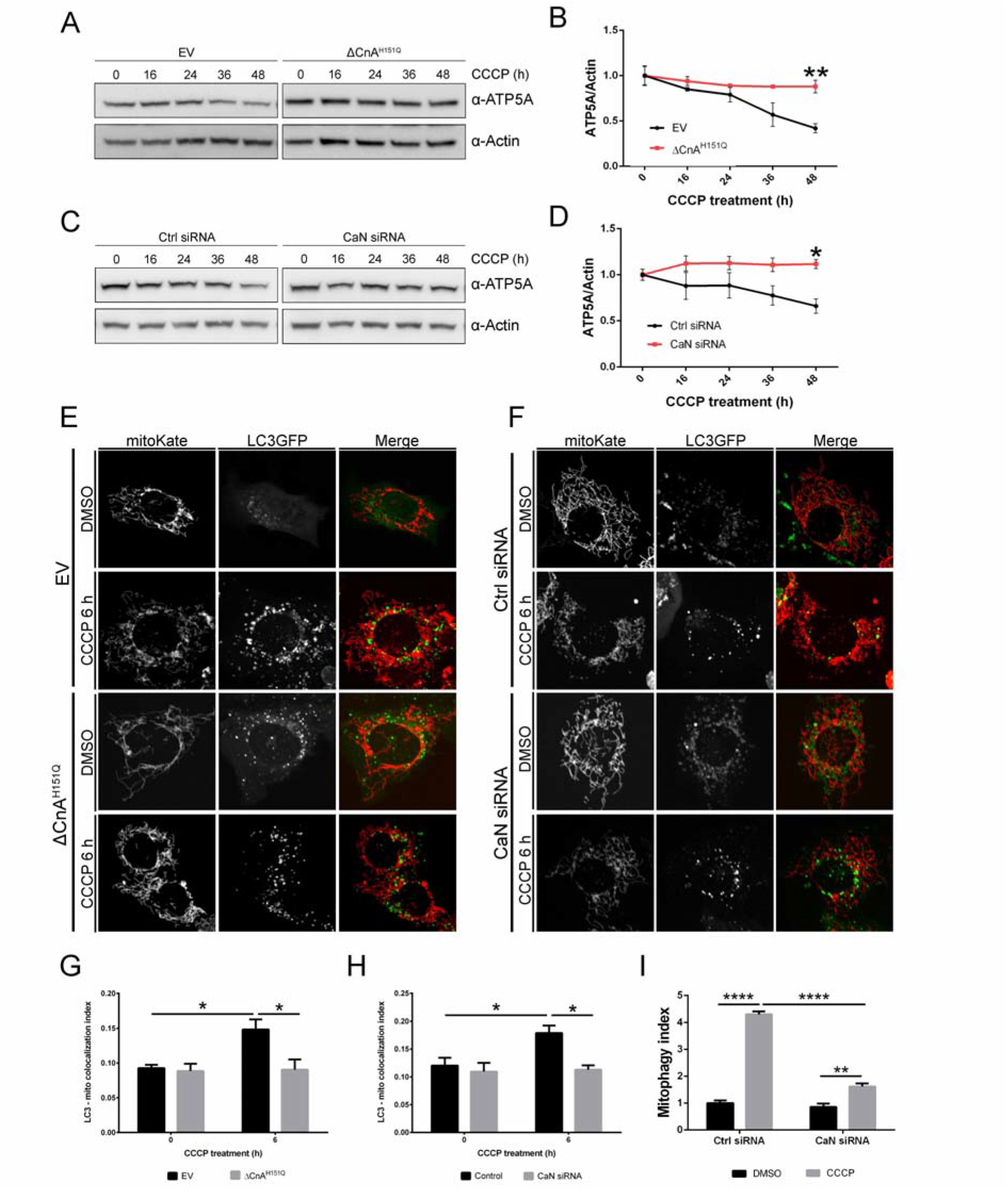
Calcineurin is required for CCCP-induced mitophagy. (A) Western blot analysis of protein lysates extracted from MEFs expressing empty vector (EV) or dominant negative CaN (ΔCnA^H151Q^). Cells were treated with 10 μM CCCP for the indicated time. (B) Quantification of (A). Line chart shows mean±SEM of ATP5A protein level normalized to Actin. Student’s t-test (n=3; p<0.01). (C) Western blot analysis of protein lysates extracted from MEFs downregulating Calcineurin and control. Cells were treated with 10μM CCCP for the indicated time. (D) Quantification of (C). Line chart shows mean±SEM of ATP5A protein level normalized to Actin. Student’s t-test (n=5; p<0.05) (E) Representative confocal images of cells transfected with LC3GFP and MitoKate plus dominant negative CaN (ΔCnA^H151Q^) or empty vector (EV). (F) Representative confocal images of cells transfected with LC3GFP and MitoKate in CaN downregulating condition and matching control. (G) Quantification of (E) using Squassh. Two-way ANOVA followed by Tukey’s multiple comparison test (n=3; p<0.05) (H) Quantification of (F) using Squassh. Two-way ANOVA followed by Tukey’s multiple comparison test (n=3; p<0.05) (I) mt-Keima analysis in CaN downregulation condition upon CCCP treatment. mt-Keima demonstrates a greater than 4-fold change in ratiometric fluorescence in CCCP treated cells, which was blunted upon CaN downregulation. Two-way ANOVA followed by Tukey’s multiple comparison test (n=3; p<0.001).

### Constitutive active Calcineurin induces an increase in basal mitophagy

We previously demonstrated that the expression of constitutive active CaN promotes Parkin translocation even in absence of CCCP treatment. To address whether this was also the case for mitophagy, we performed mt-Keima assay in Parkin expressing MEFs transiently transfected with constitutively active CaN. Representative image of mt-Keima acquired at confocal microscope clearly showed increased number of red fluorescence in control cells expressing constitutively active CaN, ΔCnA (Figure 7A). In these cells basal mitophagy (i.e. untreated cells) was increased (Figure 7B). Qualitative electron microscopy demonstrated an increase in autophagosomes and autolysosomes with mitochondrial like-structures inside in this condition (Figure 7C-D).

**Figure 7:**
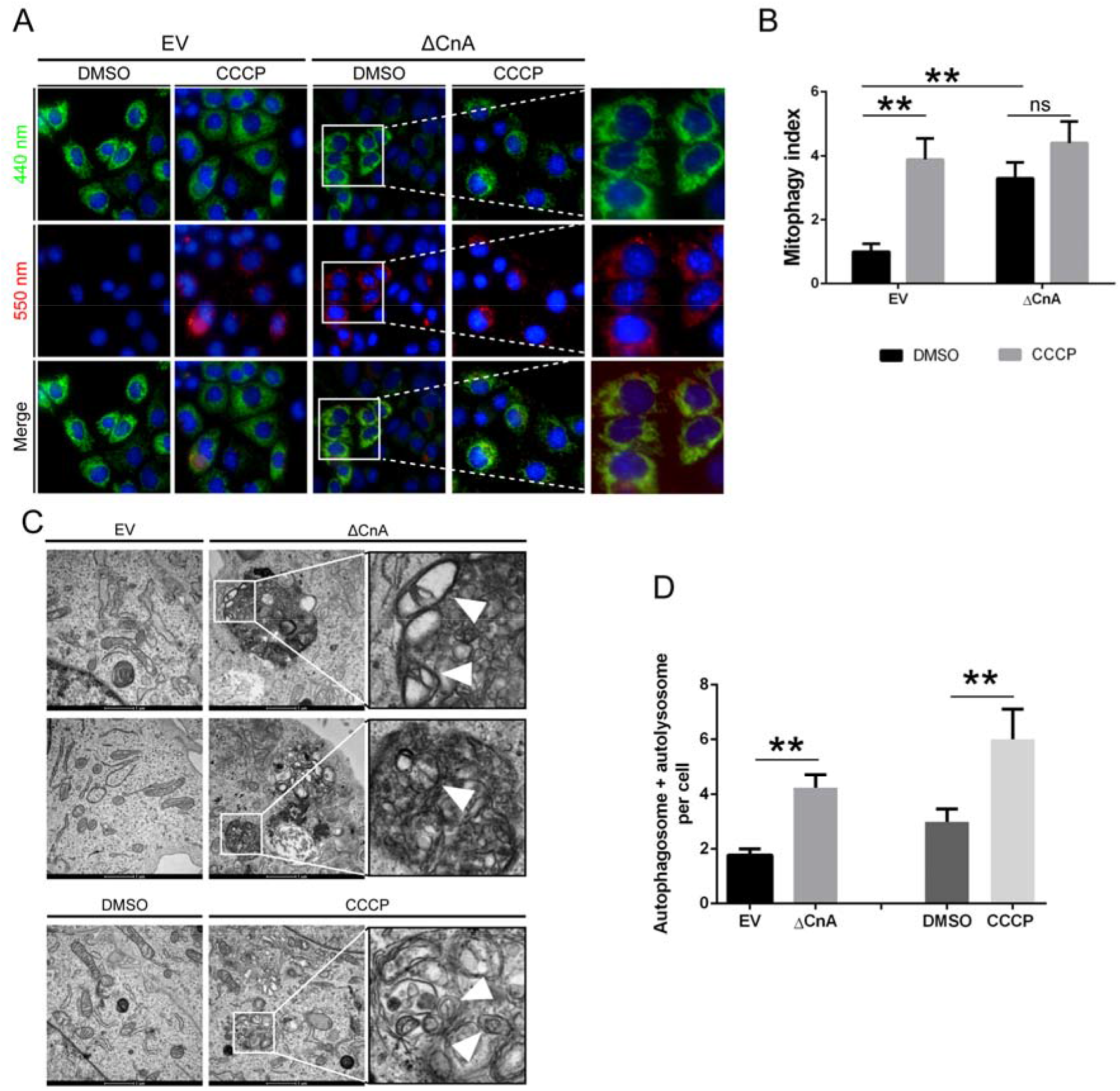
Calcineurin is required for CCCP-induced mitophagy. (A) Representative image of mt-Keima expressing cells acquired on confocal microscope Operetta High-Content Imaging system. MEFs were transfected with constitutive active CaN (ΔCnA) or empty vector (EV). (B) mt-Keima analysis in MEFs expressing constitutive active CaN (ΔCnA) and relative control (EV). Two-way ANOVA followed by Tukey’s multiple comparison test (n=6; p<0.01). (C) Representative electron microscope images showing the presence of autophagosomes with mitochondrial-like structures inside in MEFs expressing constitutive active CaN (ΔCnA) and control (EV). 24hrs CCCP-treated cells were used as positive control. Mitochondrial-like structures were identified as described in Chakraborty et al.^124^ (D) Bar graph represents mean ± SEM of the number of autophagosomes and autolysosomes per cell. At least 60 cells per biological replicate were analyzed from each group. Student’s t-test (n=3-4; p<0.01).

### Calcineurin requires PINK1 and mitochondrial fission to promote mitophagy in MEFs

While investigating Parkin translocation in PINK1 KO background, we came across the unexpected finding that expression of constitutive active CaN triggered Parkin translocation in the absence of PINK1 (Figure 3). Parkin translocation induced by CaN is independent of Drp1 activity and Drp1 mitochondrial recruitment to promote mitochondrial fission (Figure 5). Based on these observations, we now want to address the potential “mitophagic” effect of CaN activation in PINK1 KO background. Importantly, expression of constitutive active CaN in PINK1 KO cells failed to promote the degradation of mitochondrial protein ATP5A (Figure 8A-B). These quantitative immunoblotting results were confirmed by mt-Keima assay (Figure 8C). Analysis of protein levels of *bona-fide* Parkin substrates^48^ revealed that the levels of TOM20, Mfn1 and VDAC were decreased when constitutive active CaN was expressed in WT and PINK1 KO background (Figure 8D-E). In this condition, we also tracked an increase in the coefficient of colocalization between Ubiquitin and TOM20 (Figure 8F-G). Importantly, in MEFs that do not stably express Parkin, expression of constitutive active CaN failed to enhance mitophagy (Supplementary figure 9) further supporting the hypothesis that CaN-induced mitophagy is mediated by Parkin and not other E3 ubiquitin ligases that are known to share common mitochondrial targets (for example MARCH5^83^ and/or Mul1^84^). Of note, the effect of CaN on basal mitophagy is not secondary to membrane potential alteration because we did not record any significant difference in mitochondrial membrane potential upon expression of constitutive active CaN (Supplementary figure 10).

**Figure 8:**
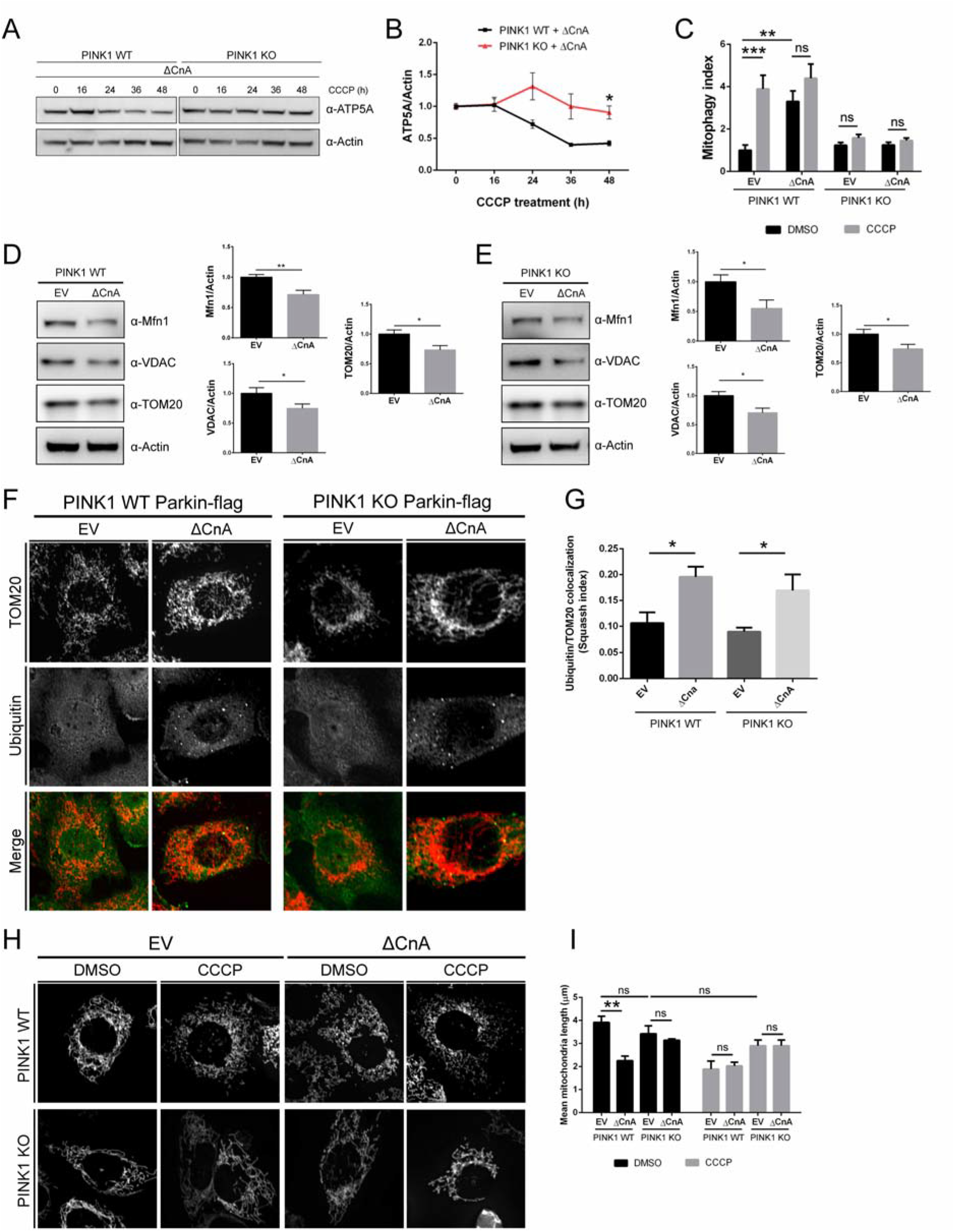
Calcineurin requires PINK1 and mitochondrial fission to induce mitophagy in MEFs. (A) Representative Western blot analysis of protein lysates extracted from PINK1 WT and KO MEFs. Cells were transfected with constitutive active CaN (ΔCnA) and after 2 days they were treated with 10 μM CCCP for the indicated time. (B) Quantification of (A). Line chart shows mean±SEM of ATP5A protein level normalized to Actin. Student’s t-test (n=5; p<0.05). (C) mt-Keima analysis of wild type (WT) and PINK1 KO MEFs. Cells were transfected with constitutive active CaN (ΔCnA) or empty vector (EV) for 2 days before being treated with DMSO or 10 μM CCCP for 3hrs, and subjected to FACS analysis. Two-way ANOVA followed by Tukey’s multiple comparison test (n=6; p<0.01). (D) Representative Western blot analysis of protein lysates extracted from PINK1 WT and KO MEFs. Cells of the indicated genotype were transfected with constitutive active CaN (ΔCnA) or empty vector (EV). Graph bars shows mean±SEM of indicated mitochondrial protein level normalized to Actin. Student’s t-test (n=8; p<0.001). (E) Representative Western blot analysis of protein lysates extracted from PINK1 WT and KO MEFs. Cells of the indicate genotype were transfected with constitutive active CaN (ΔCnA) or empty vector (EV). Graph bars shows mean±SEM of indicated mitochondrial protein level normalized to Actin. Student’s t-test (n=11; p<0.05). (F) Representative confocal images of cells of the indicated genotype transfected with constitutive active CaN (ΔCnA) or empty vector (EV). Cells were pretreated with proteasome inhibitor MG132 before being fixed, permeabilized and incubated with the indicated primary antibodies and corresponding fluorophore-conjugated secondary antibodies. (G) Quantification of (F) using Squassh. Two-way ANOVA followed by Tukey’s multiple comparison test (n=4; p<0.05). (H) Representative confocal images of cells of the indicated genotype transfected with mito-YFP and constitutive active CaN (ΔCnA) or empty vector (EV), treated with DMSO or 10μM CCCP for 3hrs. (I) Quantification of (H) using Squassh. The graph bar shows mean ± SEM of mean object length of each image for at least ≥ 70 images per biological replicate. Two-way ANOVA followed by Tukey’s multiple comparison test (n=3; p<0.05).

Because PINK1 favours Drp1 activation via PKA displacement^51^ and directly phosphorylates Drp1 to promote Drp1-dependent fission^53^, it is possible that in PINK1 KO MEFs, mitochondria simply do not fragment to allow efficient mitochondrial autophagy. To address this hypothesis, we investigated by confocal live imaging mitochondrial length of WT and PINK1 KO cells transiently expressing constitutive active CaN (ΔCnA) and subjected to 3hrs CCCP. Expression of constitutive active CaN promotes mitochondrial fragmentation in WT cells, which was not exacerbated in presence of CCCP. On the contrary, the mitochondrial network of PINK1 KO cells did not fragment when constitutive active CaN was expressed (Figure 8H-I). In addition, although PINK1 KO cells did not seem to have more elongated mitochondria than WT cells under basal condition, mitochondria did not fragment when treated with CCCP, indicating impaired fission in this condition (Figure 8H-I).

Thus, in PINK1 KO MEFs expressing constitutive active CaN, mitochondria do not fragment and cannot be efficiently eliminated by mitochondrial autophagy. In this condition, Parkin appears to be on ubiquitinated mitochondria, and protein levels of its targets are decreased.

### Constitutive active Calcineurin corrects locomotor ability and mitochondrial function of PINK1 KO flies

To evaluate the physiological relevance of our finding *in vivo*, we turned to a well-established *drosophila* model of mitochondrial dysfunction that was successfully used in many studies before: the PINK1 KO flies^85–92^. At the systemic level, PINK1 KO flies show characteristic locomotor defects in flight and climbing ability, degeneration of muscle fibers of the thorax and reduced lifespan. The phenotype of PINK1 KO flies can be rescued by overexpression of Parkin, whereas Parkin KO phenotype, which phenocopies PINK1 KO, cannot be rescued by PINK1 overexpression, and combined mutation of PINK1 and Parkin genes does not exacerbate the phenotype^86–88^. These evidences have been interpreted as PINK1 and Parkin interacting functionally in a linear pathway with PINK1 operating upstream of Parkin^86–88^. With that in mind and to assess the role of CaN in this pathway, we evaluated the effect of CaN inhibition and activation *in vivo* in PINK1 KO flies. To do so, we used a locomotor assay in which 10 flies for each strain were collected in a vertical plastic tube positioned with a line drawn at 6 cm from the bottom of the tube and under a light source. After tapping the flies to the bottom of the tube, the flies that successfully climbed above the mark after 10 seconds were counted (Figure 9A). As already reported^87,88^, PINK1 KO flies (PINK1B9) performed poorly in the climbing assay compared to wild type (Figure 9B), and Parkin overexpression (Parkin OE) rescued PINK1 KO climbing defects (Figure 9B). Feeding PINK1 KO flies with CaN specific inhibitor FK506 diminished the rescuing effect of Parkin overexpression on climbing performance (Figure 9C), while FK506 administration alone did not show any adverse effects on the climbing ability of PINK1 KO flies (Figure 9D). Importantly, expression of CaN constitutive active (CanA-14F)^72^ in PINK1 KO background rescued PINK1 KO climbing defects (Figure 9D), an effect that was suppressed by feeding flies with CaN inhibition FK506 (Figure 9D). Moreover, expression of constitutive active CaN enhanced basal mitophagy in WT and PINK1 KO flies (Figure 9E-F), and rescued PINK1 KO mitochondrial respiratory defects (Figure 9G-H).

**Figure 9:**
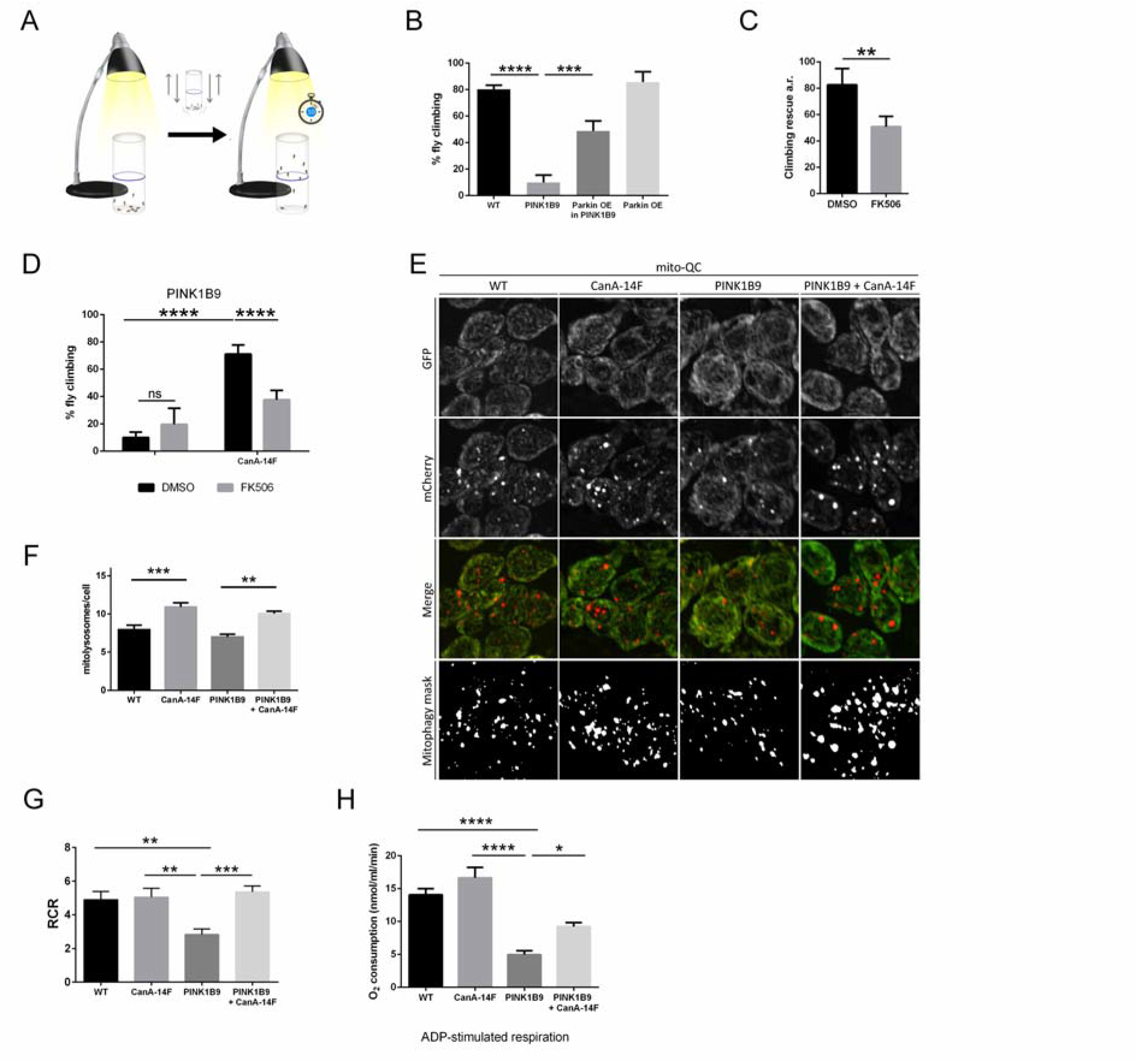
Constitutive active Calcineurin corrects locomotor ability and mitochondrial function of PINK1 KO flies. (A) Schematic representation of the climbing assay. 10 flies were put into a tube in a dark room. A light was put on the top of the tube. After tapping the flies at the bottom of the tube, the number of flies that successfully climbed above the 6-cm mark after 10 seconds was recorded. (B) Graph bar shows mean±SEM of the climbing performance of flies of the indicated genotype. One-way ANOVA followed by Sidak’s multiple comparison test (n=5; p≤ 0.001). (C) Graph bar shows mean±SEM of the climbing rescue (arbitrary unit) of PINK1 KO flies (PINK1B9) overexpressing Parkin and treated as indicated for 48hrs with DMSO or FK506. Student’s t-test (n=5; p≤ 0.01). (D) Graph bar shows mean±SEM of the climbing performance of PINK1 KO flies (PINK1B9) upon expression of constitutive active calcineurin (CanA-14F), and treated as indicated for 48hrs with DMSO or FK506. Two-way ANOVA followed by Sidak’s multiple comparison test (n=5; p≤ 0.001). (E) Representative confocal microscopy images of neurons in the ventral nerve chord of third instar larvae. Fluorescence corresponding to neutral pH (GFP) and acidic pH (mCherry) are shown. Mitolysosomes are counted as GFP-negative/mCherry-positive (red-only) puncta. The mitophagy mask generated by the mito-QC Counter allows visualization of the quantified mitophagy areas^123^. (F) Quantification of total number of mitolysosomes per cell using the mito-QC Counter^123^ (Fiji). At least 40 cells were analyzed per animal. One way ANOVA followed by Sidak’s test for multiple comparisons (n=6; p ≤ 0.01). (G) Quantitative analysis of respiratory fitness of mitochondria from adult flies of the indicated genotype. Graph shows respiratory control ratio (RCR) calculated as described in Materials and Methods. One-way ANOVA followed by Sidak’s test for multiple comparison test (n=10-11, p≤ 0.01). (H) Quantitative analysis of respiratory fitness of mitochondria from adult flies of the indicated genotype. Graph bar shows ADP-stimulated respiration calculated as described in Materials and Methods. One-way ANOVA followed by Sidak’s test for multiple comparison test (n=10-11, p≤ 0.01).

## Discussion

Mitophagy, a selective kind of autophagy during which defective mitochondria are recognized and degraded, depends on Serine/threonine kinase PINK1 and E3 ubiquitin ligase Parkin^93^, two genes which mutations have been linked to the onset of an autosomal recessive juvenile parkinsonism^9,94^. Previous studies independently showed that during stress-induced mitophagy, the E3 ubiquitin ligase Parkin translocates in a PINK1-dependent manner to depolarized mitochondria^33,38,46^. In this process, kinase PINK1 phosphorylates Ubiquitin^43^, Parkin^95^ and its targets^49,96^, and promotes mitochondrial Parkin translocation^38^ and Parkin activity^43,97^. On depolarized mitochondria, Parkin ubiquitinates mitochondrial pro-fusion proteins Mitofusin (Mfn)^38,47,48,98,99^ leading to its chaperone p97/VCP-mediated retrotranslocation for proteosomal degradation^47^. In addition, Parkin ubiquitinates the mitochondrial protein translocase TOM20, mitochondrial VDAC/Porin and Fis1^48^, and it also promotes the degradation of Miro^96^, a protein that couples mitochondria to microtubules. Selected mitochondria are therefore deprived of their pro-fusion protein Mfn, they are isolated from the mitochondrial network, and driven to autophagic degradation via autophagy adaptors like p62 (SQSTM1)^100^, HDAC6^101^, Optineurin and NDP52^102^. As shown in many studies, CCCP treatment is one of the most consolidated stimuli to promote Parkin translocation and stress-induced mitophagy. CCCP triggers mitochondrial depolarization by transporting protons inside the mitochondrial matrix, and induces a transient increase of cytosolic Ca^2+^ influx^57^ that leads to CaN activation^52^; CaN activation promotes Drp1 translocation to mitochondria to induce mitochondrial fission^52^, and the transcription of autophagy and lysosomal genes via TFEB^60^. In perfect agreement with this coordinated set of events, we found that activated CaN interacts with the mitophagic protein Parkin, and promotes Parkin mitochondrial recruitment. Importantly, the effect of CaN activation on mitochondrial recruitment of Parkin is PINK1-independent, and does not affect mitochondrial membrane potential. Parkin-dependent mitophagy has been described in the absence of mitochondrial membrane potential depolarization, following proteotoxic stress^55,103^ or upon specific induction of mitochondrial Ca^2+^ oscillation^104^. In this scenario, PINK1 is presumably not stabilized on the outer mitochondrial membrane to promote Parkin recruitment because it is imported by a functional mitochondrial import machinery, and rapidly degraded^39^. Also, PINK1-independent mechanisms of Parkin recruitment have been previously described^105^. These studies suggest that Parkin recruitment and Parkin-dependent mitophagy can be activated upon stimuli, which do not necessarily culminate in mitochondrial damage or mitochondrial ROS production, and more importantly that alternative PINK1-independent mechanisms of Parkin recruitment and activation can be predicted. It is possible that additional kinases other than PINK1 can phosphorylate Parkin, Ubiquitin, and prime mitochondria for Parkin recruitment and mitophagy. Parkin is a heavily phosphorylated protein, and several kinases have been reported to phosphorylate Parkin (for example AMPKA1^106^, CDK5^107^, PLK1^108^, ULK1^109^ and GSK3β^110^). It is possible that one of these kinases or another kinase yet to be identified are responsible for Ubiquitin phosphorylation, mitochondrial priming, and Parkin activation and recruitment, or that phospho-ubiquitin independent mechanisms of Parkin activation, such as Parkin neddylation^110,111^, are triggered by CaN activation. Undoubtedly, the mechanism underlying activation and mitochondrial recruitment of Parkin is not completely unravelled, and remains fertile ground for further debate. Interestingly, although Parkin is efficiently recruited to mitochondria in PINK1 KO cells expressing constitutive active CaN, this is not sufficient to promote mitophagy *in vitro*, presumably because in MEFs, Drp1-dependent mitochondrial fission is required to allow efficient engulfment of damaged mitochondria. As shown in many *in vitro* studies, the autophagic engulfment of mitochondria need to be paralleled by a tight control of mitochondrial size, and mitochondria need to fragment prior to mitophagy^59^. Because PINK1 inhibits PKA-mediated phospho-inhibition of Drp1^51^ and it directly phosphorylates Drp1 on S616 to regulate mitochondrial dynamic^53^, PINK1 expression is required in MEFs to release Drp1 inhibition and promote its mitochondrial recruitment to initiate mitochondrial fragmentation. Following mitochondria damage, PINK1 becomes active, disrupts the AKAP1-PKA and phosphorylates Drp1 to promote Drp1-dependent fission^51^, which is required to execute mitophagy. Thus in MEFs, expression of CaN does not promote mitophagy in the absence of PINK1, although Parkin is efficiently recruited to mitochondria, and protein levels of its mitochondrial targets TOM20, VDAC and Mfn1 are decreased.

At the systemic level, CaN activation in PINK1 KO flies rescues the characteristic locomotor defects in climbing ability and mitochondrial dysfunction of these flies. Importantly, in the brain of the fly larvae, CaN expression enhances basal mitophagy even in the absence of PINK1 expression. This result highlights a fundamental difference between CaN-dependent mitophagy in MEFs and neuronal cells of the fly head: in MEFs, PINK1 expression seems to be an absolute prerequisite for mitophagy execution, presumably because PINK1 is required to promote Drp1-dependent fission. It is possible that in the neurons of the PINK1 KO fly brain, mitophagic events predominately occur in a piece meal fashion, by selective targeting portions of mitochondria for turnover via mitochondria derived vesicles (MDV)^112,113^, a process that does not seem to require Drp1-dependent fission^112^. How mitochondrial autophagy is differentially executed in specific cell type and in the complexity of the whole organism remains poorly understood, and open to new discoveries.

As observed by others before us^91^, we did not see defective mitophagy in the brain of PINK1 KO flies, raising questions regarding the physiological relevance of PINK1 activation *in vivo* during mitophagy. One possible explanation for this lack of effect might be that PINK1-independent mitophagic pathways are activated during development to compensate for PINK1 loss. Recent publications provide hints toward this hypothesis, and suggest that the mitochondrial quality control pathways are intertwined so that activation of one specific pathway can compensate for loss of another^84,114,115^. Nevertheless, PINK1 KO flies do present a very obvious mitochondrial phenotype: mitochondrial ultrastructure is deranged, Complex I activity is compromised, and mitochondrial respiration is defective^85–87,90,116^. These conditions increase the risk of oxidative stress and inflammation deriving from cytosolic release of mitochondrial DAMPs and exposure of mitochondrial antigens^117^. In this scenario, enhancement of PINK1-independent mitophagy can promote elimination of dysfunctional mitochondria, prevent ROS builds up, and mitigate inflammation. While we cannot exclude the possibility that CaN promotes pro-survival and neuroprotective pathways via functions that are independent of Parkin-driven mitophagy, our *in vivo* results fully support the hypothesis that the rescue depends at least in part on an amelioration of mitochondrial quality control, which in the fly head it occurs in the absence of PINK1.

In conclusion, this study highlights an unprecedented role for Ca^2+^-dependent phosphatase CaN in the regulation of Parkin translocation and mitophagy. A transient increase of cytosolic Ca^2+^ influx promotes activation of CaN, which interacts with Parkin and induces Parkin mitochondrial recruitment. In parallel, CaN activates the transcription of autophagy and lysosomal genes via TFEB^60^, and allows Drp1 mitochondrial recruitment and mitochondrial fission to execute mitophagy (Figure 10).

**Figure 10:**
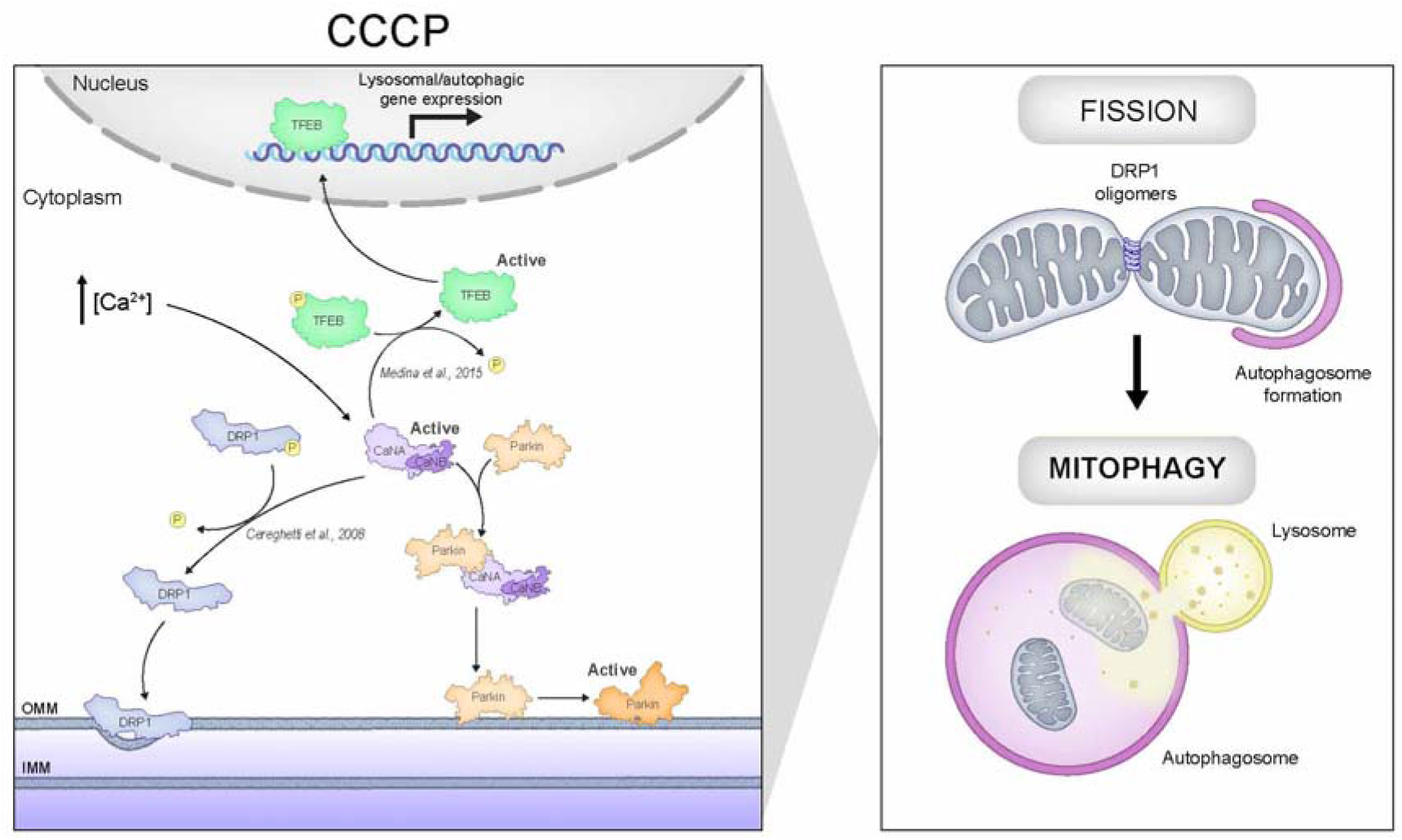
Schematic representation of the pathway regulating Parkin translocation and mitophagy induced by Calcineurin. Mitochondrial membrane potential drives PINK1 import into healthy mitochondria through the TOM and TIM complexes. Once on the IMM, PINK1 gets cleaved by MPP and PARL and eventually degraded by the ubiquitin-proteasome system^46,125^. In this scenario, CaN is not active and Parkin is kept in the cytosol. Mitochondria depolarization induced by CCCP is followed by cytosolic Ca^2+^ rise^57,125^, which activates CaN^126^. Activated CaN promotes mitochondrial recruitment of Drp1 and Drp1-dependent mitochondrial fission by dephosphorylating Drp1^52^. Activated CaN also promotes dephosphorylation of TFEB to induce its activation and the expression of autophagy and lysosomal genes^60^. In parallel, CaN interacts with Parkin and promotes Parkin translocation to mitochondria in a PINK1-independent fashion. The interaction between CaN and Parkin might be direct or via a binding partner.

## Materials and Methods

### Cells

Mouse embryonic fibroblast cells (MEFs) and Human embryonic kidney 293T (HEK293T) were cultured in Dulbecco’s modified Eagle medium (DMEM) (Thermo Fisher Scientific) supplemented with 10% Fetal Bovine Serum (FBS), 2 mM L-glutamine, 1% MEM non-essential amino acids solution, 50 U/ml penicillin and 50 μg/ml streptomycin (Thermo Fisher Scientific),and incubated at 37°C, in a humidified 5% CO2 atmosphere. PINK1 wildtype and knock-out cells were kindly provided by Prof. Francesco Cecconi lab (Danish Cancer Society, Denmark) and Prof. Juan Pedro Bolaños lab (Universidad de Salamanca, Spain).

Transfection was performed using Transfectin™ Lipid Reagent (BIO-RAD) following manufacturer instruction. 4-6 hours after transfection the medium was changed and cells were processed for the indicated experiment 24/48 hours after. This protocol has been used both for confocal microscope analysis and for protein assays. Alternatively, cells were transiently transfected with plasmid DNA using PEI (Polysciences, 24765) according to manufacturer’s instructions. Cells were silenced with Calcineurin siRNA oligo duplex (OriGene, SR416619) by direct transfection, using Transfectin (BIO-RAD) according to the protocol of the manufacturer. siRNA transfected cells were collected after 48hrs.

The following drugs were used: CCCP 10 μM (Sigma-Aldrich), Rotenone 2μM (Sigma-Aldrich), Oligomycin 1 mg/ml (Sigma-Aldrich), Antimycin A 2.5 μM (Sigma-Aldrich), FK506 0.6μM (Sigma-Aldrich).

### Constructs and Molecular Biology

mCherry-Parkin and HA-Ubiquitin plasmids were obtained from Addgene. Site directed mutagenesis, using QuickChange II XL kit (Agilent) and the following primers were used to generate a point mutation on Serine 65 in Parkin (S65E): F-MutpkSer65E (5’-GAC CTG GAT CAG CAG GCC ATT GTT CAC ATT GT-3’) and R-MutpkSer65E (5’-ACA ATG TGA ACA ATG GCC TGC TGA TCC AGG TC-3’). The same protocol was used for Ubiquitin point mutation at Serine65, and the following primers were used: F-MutUbSer65E (5’-ATC CAG AAG GAG GAG ACC CTG CAC CT-3’) and R-MutUbSer65E (5’-AGG TGC AGG GTC TCC TCC TTC TGG AT-3’). These constructs were named Parkin S65E and Ub-S65E.

Flag-tagged Parkin was inserted into pMSCV vector by using the pCR®8/GW/TOPO^®^ TA Cloning^®^ Kit (Thermo Fisher Scientific). To perform the pCR^®^8/GW/TOPO^®^ cloning, Flag-Parkin construct was PCR amplified from pEYFP-C1-Parkin vector (available in the lab) using the following primers: Parkin-forward-BglII-Flag (5’ -AGCT AGATCT ATG GAT TAC AAG GAT GAC GAC GAT AAG ATG ATA GTG TTT GTC AGG-3’) and EYFP-reverse (5’ -ACC ATG GTG AGC AAG GGC GAG-3’).

pcDNA3.1-ΔCnA^H151Q^ (ΔCnA^H151Q^), pDCR-HA-ΔCnA (ΔCnA), pDCR-CnB, Drp1-YFP and variants,, and mito-YFP were plasmids already available in the lab and described in^52,118,119^. pLVX-Puro-mitoKeima was kindly provided by Prof. Finkel (Center for Molecular Medicine, National Heart Lung and Blood Institute, NIH Bethesda, USA).

GFP-LAMP1 and GFP-Rab5 were purchased from Addgene.

### Production of lentiviral particles and infection

pLVX-Puro-mitoKeima was kindly provided by Toren Finkel (Center for Molecular Medicine, National Heart, Lung, and Blood Institute, NIH, Bethesda, MD, USA). HEK293T cells were seeded onto 100 mm-diameter tissue culture Petri dishes. 24 h after plating, cells were cotransfected using PEI with LV-mitokeima and the packaging plasmids pMDLg/pRRE, pRSV-Rev, pMD2.G. After 8 h, the transfection medium was replaced with fresh culture medium. 2 million MEFs were seeded onto 60 mm Petri dishes. After 48 h, the cells medium was collected and mixed with 6μg/ml of Polybrene (Sigma-Aldrich). Receiver MEFs were infected for 36 h, before changing the medium. All procedures for the production and use of lentiviral particles were performed in a biosafety level-2 (P2) environment.

### Retroviral infection and generation of Parkin-flag stable cell lines

To generate Parkin-flag retroviruses, HEK 293T cells were plated at 2 M cells/100-mm-diameter tissue culture dishes and transfected with the pN8E VSGV, Gag-pol packaging vectors and the retroviral vector (empty or containing Parkin-flag) by PEI (Polyscience) direct transfection. At 5 h post-transfection, the medium was replaced with fresh DMEM containing 10% FBS, and cells were grown for an additional 24 h, before the transfection was repeated. After 24 h, the conditioned medium containing recombinant retroviruses was collected and filtered through 0.45 μm-pore-size filters. Samples of these supernatants were applied immediately to MEFs cells, which had been plated 18 h before infection at a density of 10^5 cells/60-mm-diameter tissue culture dishes. Polybrene (Polyscience) was added to a final concentration of 6 μg/ml, and the supernatants were incubated with the cells for 24 h. After infection, the cells were placed in fresh growth medium and cultured in DMEM culture medium. Selection with 200 μg of Hygromycin B/ml (Sigma-Aldrich) was initiated 24 h after infection. After about 15 days, cells were expanded.

### Immunofluorescence (IF)

For Ubiquitin-TOM20 immunostaining, MEFs transiently expressing constitutively active CaN or corresponding empty vector were pretreated with 50 μM proteasome inhibitor Mg132 (Sigma-Aldrich) for 30 min. Cells were then fixed with 4% PFA in PBS for 15 min at room temperature, permeabilized with 0,1% Triton X-100 in PBS, and blocked with BSA 4% in PBS supplemented with 0,05% Tween20. Cells were incubated overnight with the following antibodies: Ubiquitin (1:100; Enzo Life Science; BML-PW8810) and TOM20 (1:200; Santa Cruz; sc-11415). Cells were washed three times in PBS supplemented with 0,05% Tween20 and subsequently incubated with the corresponding Alexa secondary antibodies (Thermo Fisher Scientific).

### Imaging

For confocal imaging experiments of Parkin localization, transfected MEFs cells were seeded onto 24 mm-round glass coverslips in 6-well culture plates. Cells were co-transfected with mCherry-Parkin construct together with mito-YFP. When indicated, cells where cotransfected with CnB, the regulatory Calcineurin (Cn) domain, plus ΔCnA (constitutively active Cn) or ΔCnAH151Q (dominant negative mutant of Cn), and/or one of the Ubiquitin constructs (Ub or UbS65E). Image analysis was performed using ImageJ. These constructs were then excited using 561nm or 488nm laser and using a *UPlanSApo 60x/1.35* objective (iMIC Andromeda). Stack of images separated by 0.2μm along the z-axis were acquired. The quantification was performed as calculation of the percentage of cells with Parkin puncta on mitochondria or through an ImageJ plugin for colocalization quantification (see following paragraph for details).

### Image analysis using Squassh

To quantify Parkin colocalization with mitochondria, we created maximum-intensity projections of z-series with 0.2 μm increments. Quantification was then performed by using ‘Squassh’ (Segmentation and QUAntification of Subcellular SHapes), a plugin compatible with the imaging processing softwares ImageJ or Fiji, freely available from http://mosaic.mpi-cbg.de/?q=downloads/imageJ. Squassh is a segmentation method that enables both colocalization and shape analyses of subcellular structures in fluorescence microscopy images^62^. For Parkin-mitochondria colocalization analysis, segmentation was performed with the minimum intensity threshold for the first channel set to 0.35, for the second to 0.15 and the regularization weight to 0.015. Among the three different colocalization coefficients (C_signal_, C_number_ and C_size_), we preferentially used C_number_. It has to be noted that Squassh based analysis is unbiased as this method is completely automated and performed by computer software.

Colocalization analysis of Ubiquitin with TOM20 was performed on single plane images using Squassh. For this analysis, segmentation was performed with the minimum intensity threshold for the first channel set to 0.2, for the second to 0.2 and the regularization weight to 0.05. As colocalization coefficient we used C_number_.

### Mitokeima mitophagy analysis by flow cytometry

Mitokeima expressing MEFs were analyzed by flow cytometry (BD FACSAria™ sorter) as previously reported^120^. Cells were analyzed with flow cytometer equipped with a 405-nm and a 561-nm laser. Cells were excited with violet laser (405 nm) with emission detected at 610±10nm with a BV605 detector and with a yellow-green laser (561 nm) with emission detected at 610±10nm by a PE-CF594 detector simultaneously.

### Immunoblotting

At the established time points, the medium was removed and MEFs washed with ice-cold PBS. After withdrawing PBS, cells were scraped off the wells using a plastic cell scraper, they were resuspended in 1,5 ml of cold PBS and they were centrifuged at 3’000g at 4 °C for 5 min. Supernatant was discarded and then the pellet was resuspended in an appropriate volume of radioimmunoprecipitation assay (RIPA) buffer (150 mM NaCl, 50 mM Tris-HCl, 1% NP-40, 0.25% Sodium Deoxycholate, 1 mM EDTA in distilled water and adjusted pH to 7.4) with freshly added protease inhibitors cocktail (PIC). Cells were kept on ice for 30 min. Lysate were cleared by centrifugation at 20’000 g for 15 min at 4 °C.

Protein concentrations of samples was determined using Pierce™ BCA Protein Assay Kit (Thermo Fisher scientific).

NuPAGE^®^ LDS Sample Buffer (Invitrogen) and 2-Mercaptoethanol (Sigma-Aldrich) were mixed to samples and proteins were then denaturated at 95°C for 15 min. Proteins were separated on ExpressPlus™ PAGE gels (GenScript) and transferred to PVDF membrane (MERCK-Millipore). Membranes were incubated with indicated antibodies and imaged with ImageQuant LAS4000. Band densiometry quantification was performed using ImageJ software. The following antibodies were used: Actin (1:5000; Chemicon; MAB1501), ATP5A (1:5000; Abcam; ab14748), Calcineurin (1:1000; BD Bioscience; 556350 and 1:1000; Abcam; ab52761), His (1: 5000; Proteintech; 66005-1), Mfn1 (1:600; Proteintech; 13798-1-AP) Parkin (1:1000, Santa Cruz; SC32282), Parkin (1:500, Abclonal; A0968), FLAG (1:3000; Cell Signalling; 2368S), TOM20 (1:2000; Santa Cruz; sc-11415), VDAC (1:1000; Abcam; ab15895). Canonical secondary antibodies used were sheep anti-mouse or donkey anti-rabbit HRP (GE Healthcare). Mouse TrueBlot^®^ ULTRA: Anti-Mouse Ig HRP (Rockland) was used for immunoprecipitation experiment.

### Thermal Stability Assay

Cells were plated onto 100 mm Petri dishes (10 M/dish). After 24hrs, transfection was performed using PEI (Polyscience) with ΔCnA and CnB plasmids and the corresponding empty vectors. After 48 h from the transfection, cells were resuspended in PBS and snap-freezed in liquid nitrogen. The solution was aliquot into a PCR strip and incubated at the indicated temperatures for 3 min. The lysates were centrifuged at 16’000 g for 30 min at 4 °C. The soluble fraction was loaded into SDS-PAGE gel.

### In vitro interaction assay

Bacterial expression Parkin construct (NM_004562.2 GI:169790968) was transformed into BL21 DE3 E. Coli. Overnight culture inoculated from fresh colony was grown in Terrific broth media containing 2% glucose and 50⍰μg/⍰ml kanamycin at 37⍰°C. The following morning overnight cultures were diluted to OD 600 0.1 and continued shaking at 37⍰°C until OD 600 reached 0.8, flasks were transferred to 4 °C, cultures were then induced with 0.2⍰mM IPTG and expression was allowed to proceed overnight at 18⍰°C. Cells were harvested by centrifugation at 6’000 g for 15 min at 4 °C. For purification of Parkin construct, bacteria were resuspended in buffer A (50⍰mM Hepes pH 8.0, 200⍰mM NaCl, 10⍰mM imidazole, 250⍰μM TCEP and EDTA-free Complete protease inhibitor tablets (Roche)) and lysed using a French Press homogenizer. The lysate was cleared at 45’000⍰g for 25⍰min at 4⍰°C and the supernatant was loaded into HisPur NiNTA Chromatography 1 ml cartridge (Thermo Fisher Scientific). The resin was washed with 50 ml of buffer A containing 20⍰mM imidazole. IMAC purification was performed on an ÄKTA Purifier FPLC system (GE Healthcare) and 20 fractions were eluted in imidazole gradient (20 mM ÷ 500 mM) for 20 min. After elution, the protein was desalted using Vivaspin™ ultrafiltration spin columns into Buffer A without imidazole. The protein was then loaded onto a HiLoad Superdex 75 16/60 size exclusion chromatography column (GE Healthcare) that had been pre-equilibrated in Hepes 30 mM, NaCl 150 mM, Glycerol 10%. Collected fractions were then concentrated and loaded on Buffer A equilibrated NeutrAvidin agarose resin (Thermo Fisher) for 40 min at 4 °C. In parallel, CaN-flag expressing cells were lysed and protein extract was incubated with Parkin-bound resin overnight at 4 °C under gentle rotation. The resin was washed three times and boiled at 95 °C in 2X Laemmli buffer (160 mM Tris-HCl, pH 6.8, 4% SDS, 0.7 M Sucrose, 4% β-mercaptoethanol) in order to elute the protein sample and subsequently resolved it in a SDS-PAGE. As negative control for the assay we used USP14-Flag, a protein that is not supposed to interact with Parkin. USP14-Flag expressing cells were lysed and protein extract was incubated with Parkin-bound resin and processed as just described. As additional negative control, we immobilized MEF2D-His^121^, a protein that is not supposed to interact with Calcineurin. CaN-Flag expressing cells were lysed and protein extract was divided in two fractions: first fraction was incubated with Parkin-bound resin, while the second fraction with MEF2D-bound resin overnight. Samples were processed as previously described.

### Proximity Ligation Assay (PLA)

For proximity ligation assay, we used a PLA kit from Sigma-Aldrich (Sigma-Aldrich DUO92014), and followed the manufacturer’s instructions. HeLa cells were grown on coverslips and transiently transfected with mCherry Parkin and PPP3CB-Flag using Trans-IT transfection reagent (Mirus Bio) for 18 h. They were then fixed with 4% paraformaldehyde (Electron Microscopy Sciences, Hatfield, Pennsylvania) for 10⍰min at room temperature. The blocking buffer (0.5% bovine serum albumin, 0.1% saponin, 50⍰mM NH_4_Cl in PBS) was then added to the cells for 20⍰min. The samples were washed in PBS and incubated overnight at 4°C with the primary antibodies in the blocking buffer. The following antibodies were used for the PLA experiments: anti-GM130 (1:700; BD Biosciences); anti-PPP3CB (Origene, 1:200); anti-GRASP65 (1:1000) and anti-Parkin (1:1000) were from Abcam; Alexa 488-, Alexa 633- and Alexa 568-conjugated secondary antibodies (1:400) were from Invitrogen. As negative controls we used anti GRASP65, which is not supposed to interact with Parkin. As positive controls, we performed the PLA by using GRASP65 and GM130. The PLA was performed according to the manufacturer instructions. Finally, Hoechst was incubated for 10 min at room temperature. The coverslips were then mounted on glass microscope slides with Mowiol4-88 (Sigma-Aldrich). Immunofluorescence samples were examined using a confocal laser microscope (Zeiss LSM700 confocal microscope system; Carl Zeiss, Gottingen, Germany) equipped with ×63 1.4 NA oil objective. Optical confocal sections were taken at 1 Airy unit, with a resolution of 512 × 512 pixels or 1.024 × 1024 pixels. PLA analysis was performed using Fiji. Images were subjected to thresholding, and the number of particles (“PLA dots”) was calculated with the Analyze Particles function. As negative controls we used anti GRASP65, which is not supposed to interact with Parkin. As positive control, we performed the PLA by using GRASP65 and GM130 (both expressed in the cis golgi network^74^).

### Co-Immunoprecipitation assay

MEFs and HEK293T cells were pretreated with 10 μM CCCP for 2 h and cell pellet was resuspended in ice-cold lysis buffer (Hepes 50 mM, Tween-20 0,1%, Triton X-100 1%, Glycerol 10% with freshly added phosphatase and protease inhibitor, pH 7.2). Cells were kept on ice for 30 min and the soluble fractions from cell lysates were isolated by centrifugation at 12,000g for 15 min. In the meantime, 50 μl of Protein-A Agarose (Roche) were washed and equilibrated in lysis buffer. Protein extract was pre-cleared in the equilibrated resin with 30 min incubation at 4 °C. Pre-cleared sample was quantified using Pierce™ BCA Protein Assay Kit (Thermo Fisher Scientific). For co-immunoprecipitations, 5 mg of lysate was incubated with Parkin antibody (2 μg; Santa Cruz 32282) or with Calcineurin antibody (2 μg; BD Bioscience 556350). Anti-mouse IgG was used in both cases as negative control (2 μg; Santa Cruz sc-2025) with constant rotation overnight at 4 °C. Then, 50 μl of Protein-A beads (Roche) was added to lysates and incubated with rotation for 2 h at 4 °C. After incubation, the resin was washed three times in ice-cold PBS and the samples were eluted in Laemmli buffer with β-mercaptoethanol at 70 °C for 15 min and loaded on 8% SDS-PAGE.

### Electron microscopy

Samples were fixed with 2.5% glutaraldehyde in 0.1M sodium cacodylate buffer pH 7.4 ON at 4°C. The samples was postfixed with 1% osmium tetroxide plus potassium ferrocyanide 1% in 0.1M sodium cacodylate buffer for 1 hour at 4°. After three water washes, samples were dehydrated in a graded ethanol series and embedded in an epoxy resin (Sigma-Aldrich). Ultrathin sections (60-70nm) were obtained with an Ultrotome V (LKB) ultramicrotome, counterstained with uranyl acetate and lead citrate and viewed with a Tecnai G2 (FEI) transmission electron microscope operating at 100 kV. Images were captured with a Veleta (Olympus Soft Imaging System) digital camera.

### Mitochondria membrane potential analysis

Mitochondria membrane potential was measured with the fluorescent dye Tetramethylrhodamine Methyl Ester (TMRM). MEFs cells were previously seeded on 24 mm round coverslips and transfected with ΔCnA + CnB plasmids or with the corresponding empty vectors. 48 hours after transfection, cells were washed with HBSS containing 20mM HEPES (pH 7.4). Cells were then incubated in HBSS with 1μM Cyclosporin H and 10nM TMRM for30 min at 37°C. Cellular fluorescence images were acquired at room temperature with UPlanSApo 60x/1.35 objective (iMIC Andromeda). Sequential images were acquired every 1 min. After 5 min, 2.5 μg/ml Oligomycin was added directly into the acquisition chamber. After 30 min, mitochondria were fully depolarized by the addition of 2μM of the protonophore carbonylcyanide-p-trifluoromethoxyphenyl hydrazone (FCCP) and recorded for further 5min. Images were analyzed using Image J software. Mitochondria network of each cell was identified as region of interest (ROI) and field not containing cells was used as background. The mean fluorescence intensity was measured for each ROI. Following background substraction, each value of TMRM fluorescence was normalized with initial fluorescence and expressed as relative TMRM intensity.

### Fly stocks and breeding conditions

Flies were raised on standard cornmeal medium and were maintained at 23° C, 70% relative humidity, on a 12 h light : 12 h dark cycle.

We used ActGal4 or nSybGal4 standard lines crossed with w1118, generous gifts from Dr. Alexander Whitworth (University of Sheffield) as controls. PINK1B9 and PK OE lines were already described before^86,87^ and were a kind gift from Dr. Alexander Whitworth. Mito-QC line was also a kind gift from Dr. Alexander Whitworth^91^. CanA-14F line was described before^122^ and was a kind gift by Dr Pascal Dijkers.

### Climbing assay

The climbing assay (negative geotaxis assay) was used to assess locomotor ability. Climbing data were obtained from groups of untreated wildtype, untreated PINK1B9, FK506-treated wildtype, and FK506-treated PINK1B9. Briefly, 10 flies for each strain were collected in a vertically-positioned plastic tube (length 12 cm; diameter 5 cm) with a line drawn at 6 cm from the bottom of the tube. Flies were gently tapped to the bottom of the tube, and the number of flies that successfully climbed above the 6-cm mark after 10 seconds was noted. Fifteen separate and consecutive trials were performed for each experiment, and the results were averaged. At least 30 flies were tested for each genotype or condition.

The number of flies that could climb unto, or above, this line within 10 or 20 seconds was recorded and expressed as percentage of total flies.

### Analysis of mitochondrial respiration in flies

The rate of mitochondrial O_2_ consumption was monitored using an Oxytherm System (Hansatech) with magnetic stirring and temperature control, maintained at 30°C. Five adult male flies per genotype were homogenized in respiration buffer (120 mM sucrose, 50 mM KCl, 20 mM Tris-HCl, 4 mM KH2PO4, 2 mM MgCl2, 1 mM EGTA, 1 g/l fatty acid-free BSA, pH 7.2) and the following additions were made: proline 10 mM, glutamate 10 mM, malate 4 mM, ADP 2.5 mM, oligomycin 2 μM, CCCP 1.25 μM, antimycin 1.25 μM. O_2_ consumption was obtained from the registered slope of the graph. Respiratory control ratio (RCR), state III versus state IV (ADP-stimulated respiration over oligomycin-administered respiration), was also determined from the registered graphs. Data from 10–11 independent experiments were averaged.

### Analysis of mitophagy in flies (Mito-QC analysis)

Brains from third instar larvae were dissected in PBS and fixed in 4% formaldehyde, pH 7, for 20 minutes, rinsed in PBS and mounted in Prolong Diamond Antifade mounting medium (Thermo Fischer Scientific). Samples were generally dissected in the morning and imaged in the afternoon of the same day. Because X chromosome nondisjunction is present in multiple balanced PINK1B9 mutant stocks, correct genotypes were determined by PCR-based genotyping of discarded tissue after dissection.

Fluorescence imaging was conducted using a confocal microscope (Andromeda iMIC spinning disc live cell microscope, TILL Photonics, 60× objective). Z-stacks were acquired at 0.2 μm-steps. Confocal images were analyzed using Fiji (ImageJ) software. The mito-QC Counter plugin was used to quantify the number of mitolysosomes, according to Montava-Garriga et al^123^.

### Statistical analysis

We used Origin 7.0 Professional or Prism 8 for statistical analysis. All data are expressed as mean ± SEM unless specified otherwise. Statistical significance was measured by an unpaired t-test or one-way or two-way ANOVA followed by *ad hoc* multiple comparison test. p-values are indicated in the figure legend. Data information: n=number of biological replicate; *P ≤ 0.05, **P ≤ 0.01, ***P ≤ 0.001, ****P ≤ 0.0001.

## Funding

This work was supported by grants from Italian Ministry of Health “Ricerca Finalizzata” [GR-2011-02351151], Rita Levi Montalcini “Brain Gain” program and Michael J. Fox RRIA 2014 [Grant ID 9795] to E.Z.

## Acknowledgements

We thank Dr Alexander Whitworth for kindly providing fly lines described in Materials and methods. We thank PF Dijkers for the CanA-14F fly line and F Cecconi who provided PINK1 wild type and KO MEFs.

## Conflict of interests

Authors declare no conflict of interests

## Data availability

We confirm that all relevant data are available from the authors

---

**Supplementary Figure 1:**
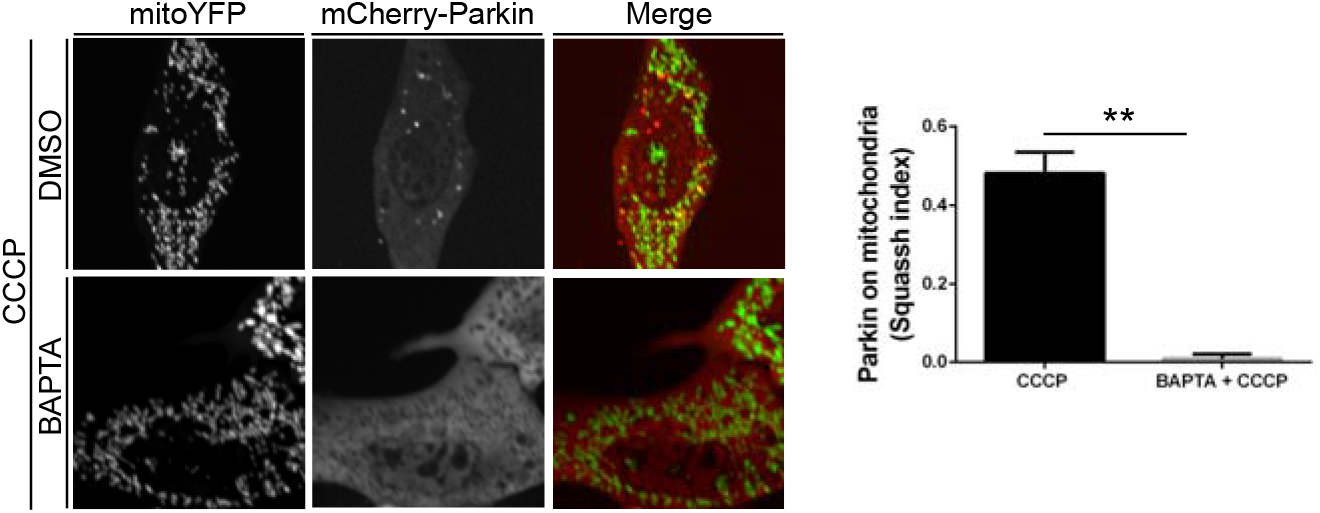
Parkin translocation to mitochondria is regulated by Calcineurin. Representative confocal images of wild type MEFs transfected with mCherry-Parkin and mito-YFP for 2 days before being treated with 40 μM BAPTA for 30 min or with 40 μM BAPTA for 30 min prior to 3 hrs 10μM CCCP treatment, as indicated. Graph bar shows mean±SEM of percentage of cells with mCherry-Parkin on mitochondria. Student’s test (n=3; p<0.01).

**Supplementary Figure 2:**
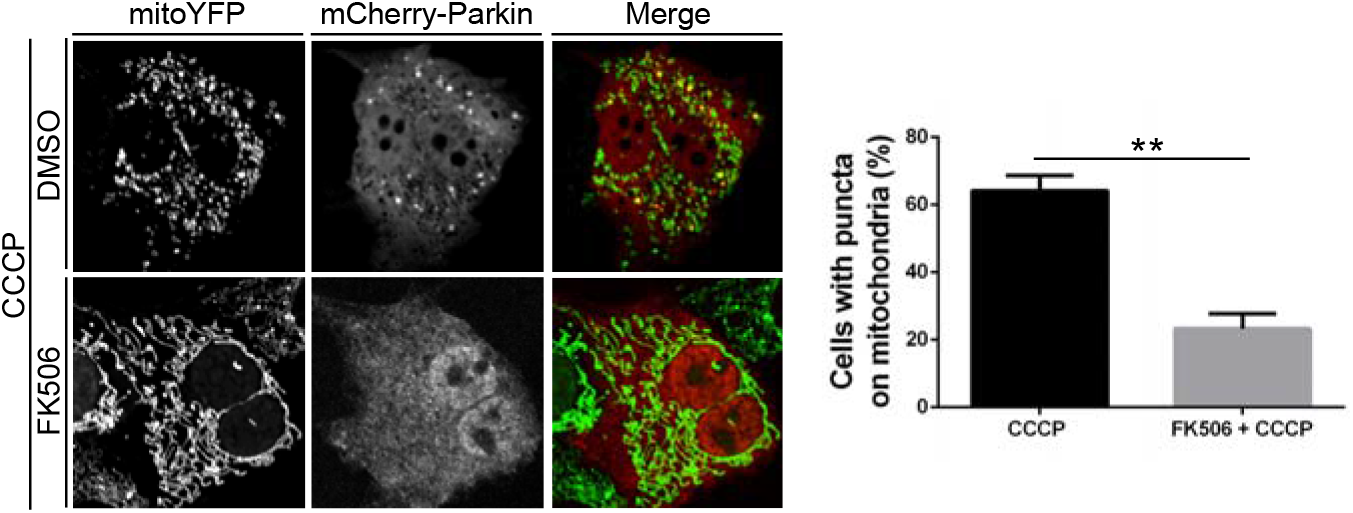
Parkin translocation to mitochondria is regulated by Calcineurin. Representative confocal images of wild type MEFs transfected with mCherry-Parkin and mito-YFP for 2 days before being treated with 0.6 μM FK506 for 30 min or with 0.6 μM FK506 for 30 min prior to 3hrs/10μM CCCP treatment, as indicated. Graph bar shows mean±SEM of percentage of cells with mCherry-Parkin on mitochondria for at least ≥ 300 cells per biological replicate. Student’s test (n=3; p<0.01).

**Supplementary Figure 3:**
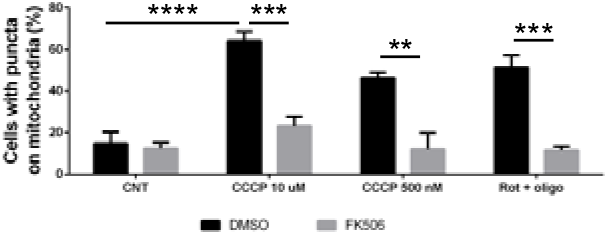
Parkin translocation to mitochondria is regulated by Calcineurin. Graph bar shows mean±SEM of percentage of cells with mCherry-Parkin on mitochondria for at least ≥ 300 cells per biological replicate. Cells were transfected with mCherry-Parkin and after 2 days they were treated as indicated. One-way ANOVA followed by Tukey’s multiple comparison test (n=3; p<0.01).

**Supplementary Figure 4:**
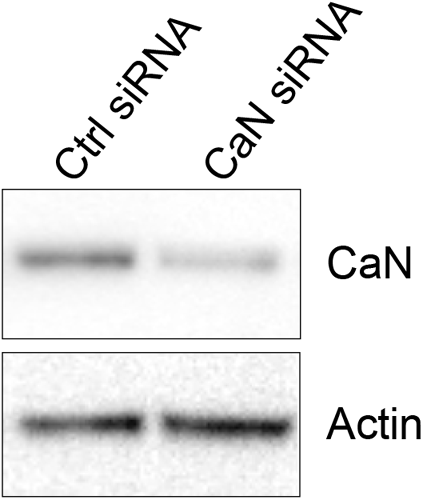
Parkin translocation to mitochondria is regulated by Calcineurin. Western blot analysis of protein lysates extracted from MEFs downregulating Calcineurin and relative control. Cells were treated with CaN siRNA and control siRNA, protein lysates were collected after 2 days and subjected to Western blotting analysis with the indicated antibodies.

**Supplementary Figure 5:**
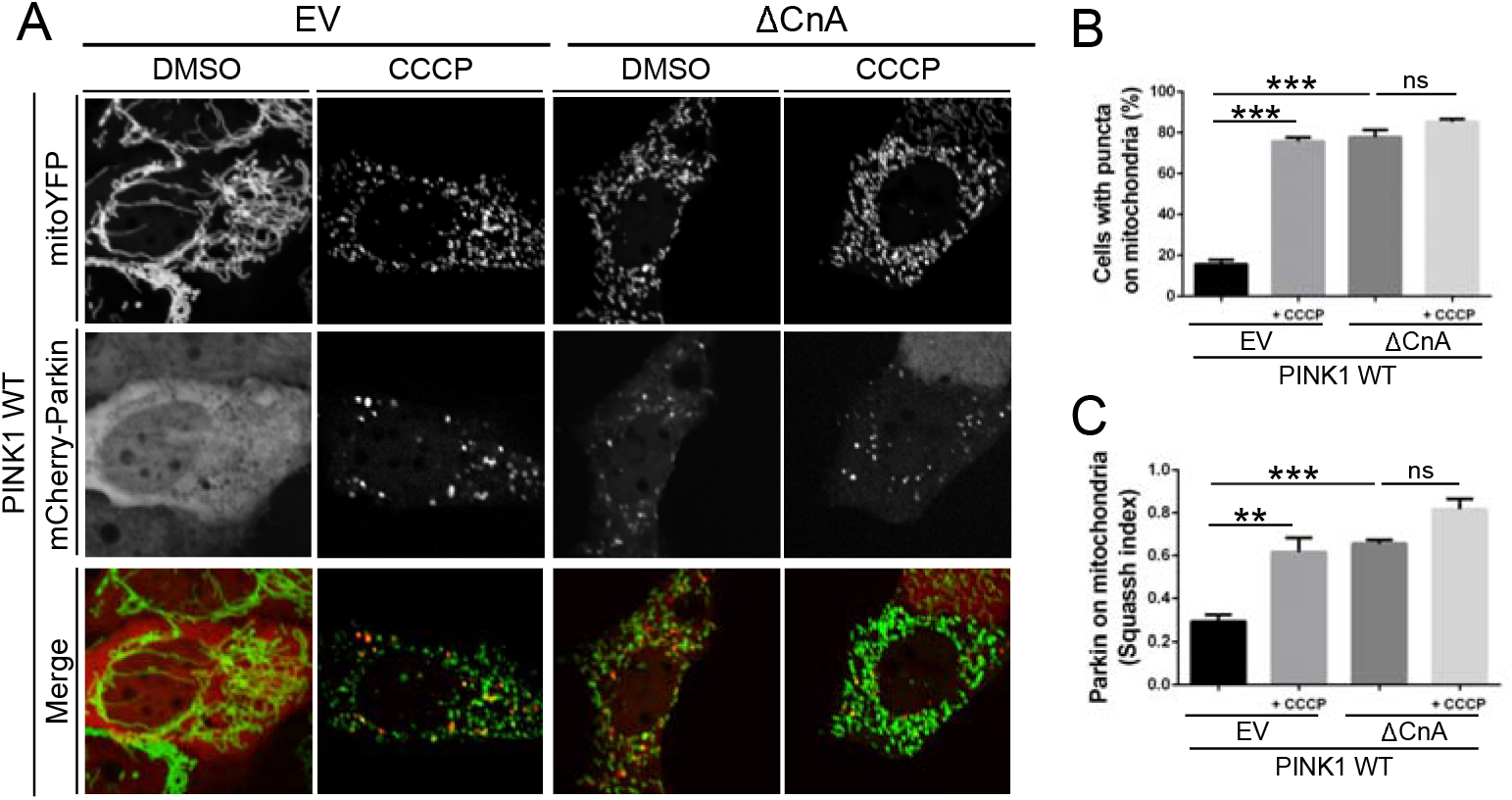
Parkin translocation to mitochondria is regulated by Calcineurin. (A) Representative confocal images of wild type MEFs transfected with mCherry-Parkin, mito-YFP and with empty vector (EV) or constitutively active CaN (ΔCnA) for 2 days before being treated with DMSO as control or 10 μM CCCP for 3hrs. (B) Quantification of A. Graph bar shows mean±SEM of percentage of cells with mCherry-Parkin on mitochondria for at least ≥ 300 cells per biological replicate. Two-way ANOVA followed by Tukey’s multiple comparison test (n=3-9; p<0.001). (C) Quantification of A by using Squassh. The graph bars show mean±SEM of Squassh colocalization coefficient for at least ≥ 50 images per biological replicate. 0=no colocalization, 1=perfect colocalization. Two-way ANOVA followed by Tukey’s multiple comparison test (n=3-4; p<0.01).

**Supplementary Figure 6:**
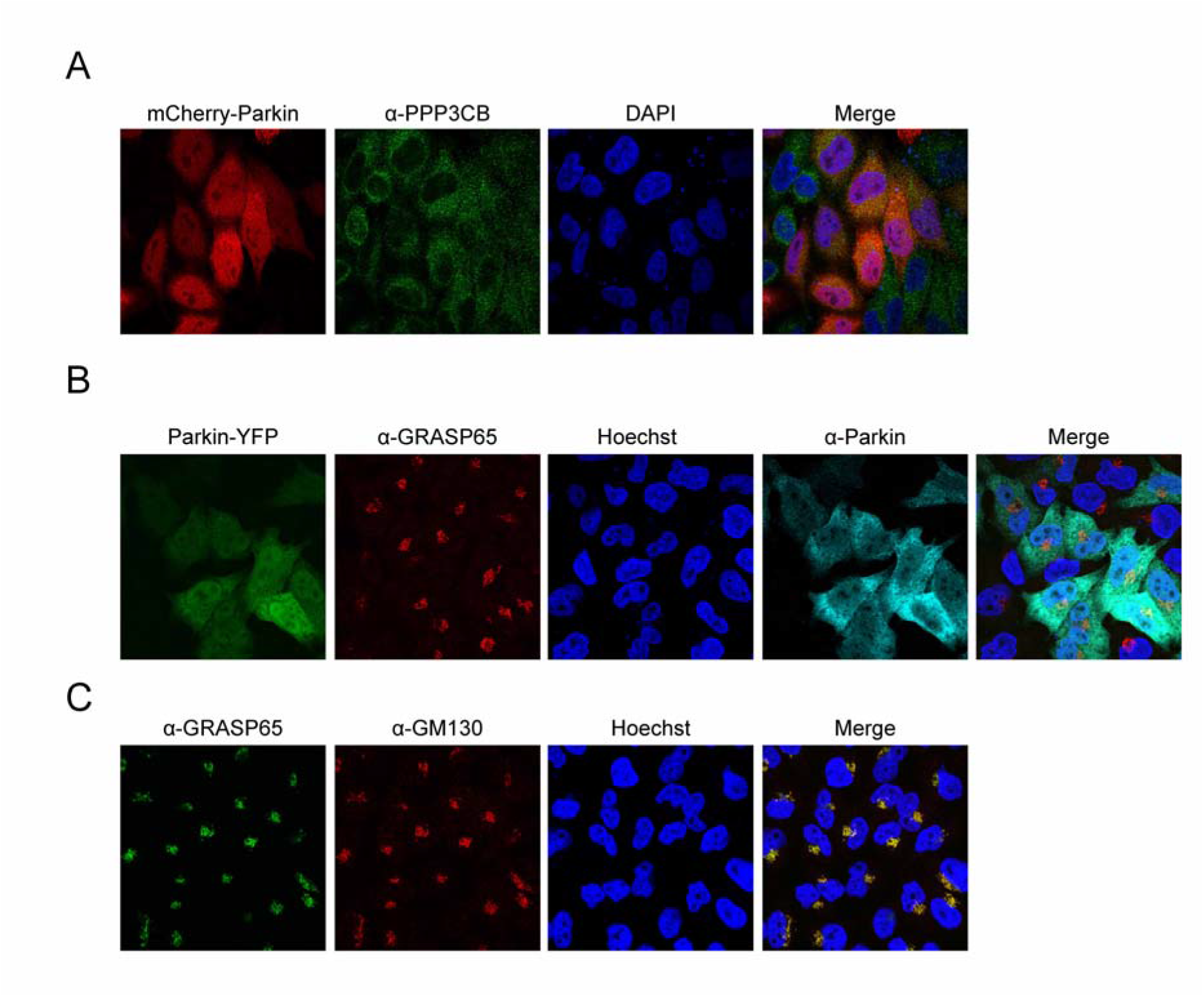
Calcineurin interacts with Parkin. (A) Representative images of HeLa cells transfected with mCherry-Parkin and probed with antibody against Calcineurin (PPP3CB). Cells were fixed, permeabilized and incubated with the indicated primary antibody, corresponding fluorophore-conjugated secondary antibody, and DAPI for nuclear staining. (B) Representative images of HeLa cells transfected with Parkin-YFP probed with antibodies against GRASP65 and Parkin. Cells were fixed, permeabilized and incubated with the indicated primary antibodies, corresponding fluorophore-conjugated secondary antibodies, and Hoechst for nuclear staining. GRASP65 is a *cis-*Golgi resident protein and does not interact with Parkin. (C) Representative images of HeLa cells probed with antibodies against GRASP65 and GM130. Cells were fixed, permeabilized and incubated with the indicated primary antibodies, corresponding fluorophore-conjugated secondary antibodies, and Hoechst for nuclear staining. GRASP65 is a peripheral membrane protein that resides in the *cis-*Golgi apparatus and interacts with GM130, which is also expressed in the *cis*-Golgi network.

**Supplementary Figure 7:**
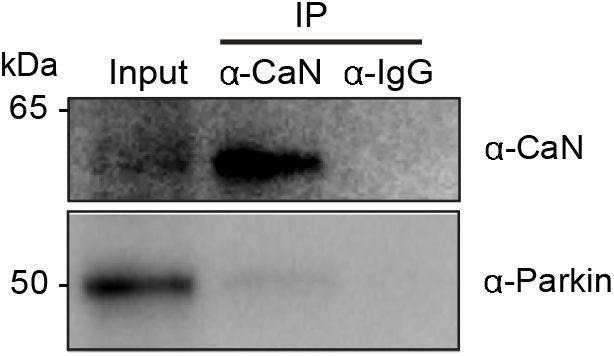
Calcineurin interacts with Parkin. HEK 293T cells were treated with CCCP-2hrs and subjected to immunoprecipitation (IP) of CaN using anti-CaN antibody. Western Blot analysis was performed with antibody anti-Parkin on the pulled down samples. Inputs represent 5% of the protein lysates and IP eluate 100% of the protein lysates.

**Supplementary Figure 8:**
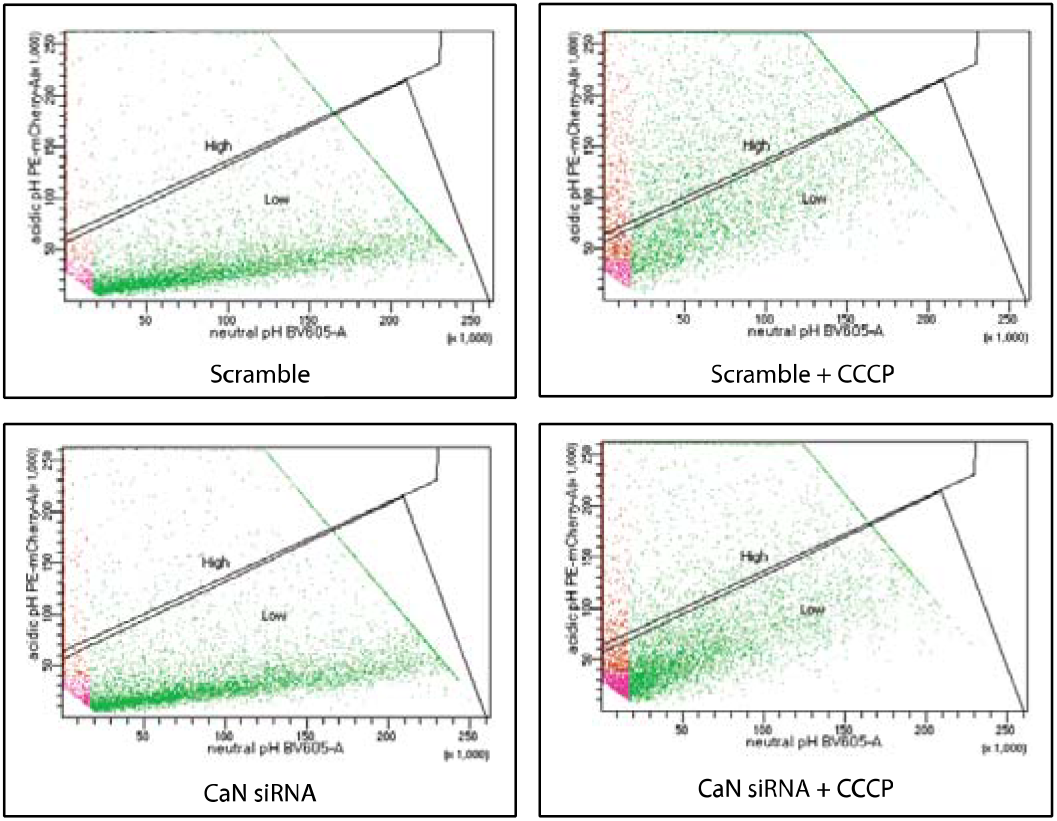
CCCP-induced mitophagy is impaired in cells downregulating Calcineurin. Representative scatterplots depicting the mean relative level of global mt-Keima signal in CaN downregulating cells and relative control.

**Supplementary Figure 9:**
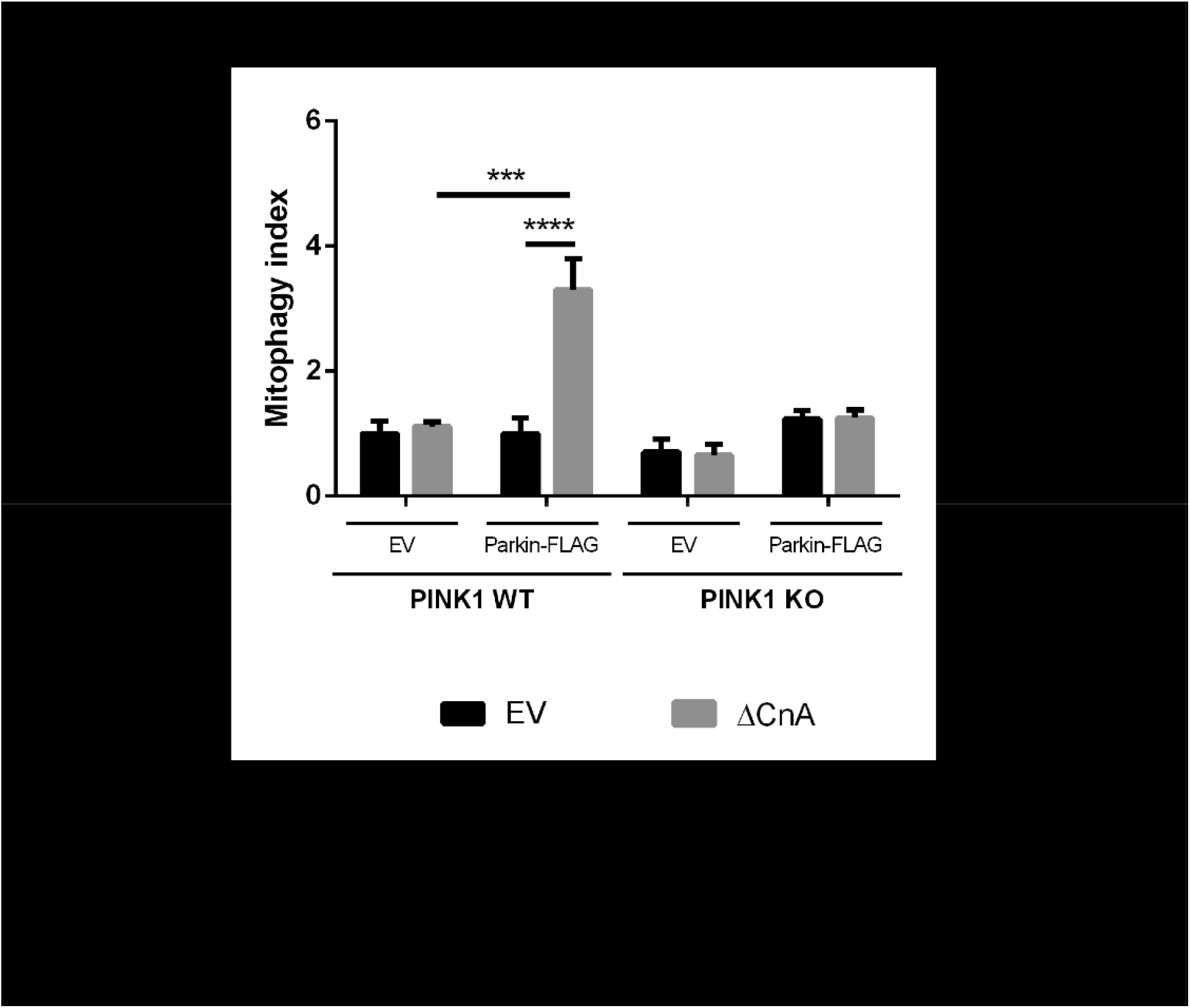
CaN-induced mitophagy is mediated by Parkin. Mt-Keima analysis in MEF cells, which do not express Parkin compared to Parkin-flag expressing cells. In PINK1 WT background, expression of constitutively active CaN failed to induce mitophagy in Parkin deficient cells, while the opposite effect was observed in stable cell line expressing Parkin-Flag. Two-way ANOVA followed by Tukey’s multiple comparison test (n≥3; p<0.001).

**Supplementary Figure 10:**
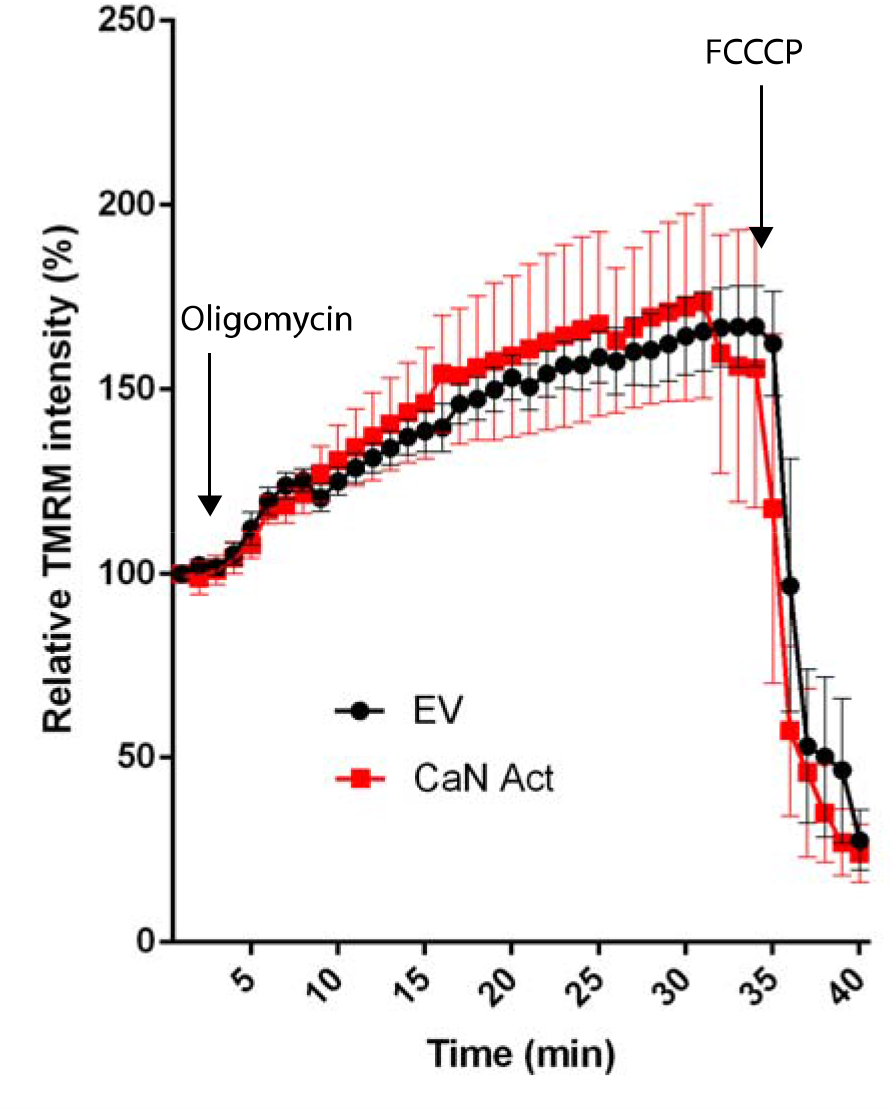
Expression of constitutive active Calcineurin does not affect mitochondria membrane potential. Analysis of mitochondria membrane potential upon expression of constitutive active CaN. MEFs cells were incubated in presence of 10nM TMRM, and images were acquired with confocal microscope. Changes of mitochondria TMRM fluorescence (expressed as the % of initial fluorescence) were followed over time.

## Notes

### Competing Interest Statement

The authors have declared no competing interest.

